# Unlocking the Secrets of NSP3: AlphaFold2-assisted Domain Determination in SARS-CoV-2 Protein

**DOI:** 10.1101/2025.01.14.632924

**Authors:** Maximilian Edich, David Briggs, Yunyun Gao, Andrea Thorn

## Abstract

Non-structural protein 3 (nsp3) is crucial for the SARS-CoV-2 infection cycle. It is the largest protein of the virus, consisting of roughly 2000 residues, and a major drug target. However, due to its size, disordered regions, and transmembrane domains, the atomic structure of the whole protein has not yet been established. Only 10 out of its 16 domains were individually determined in experiments.

Here, we demonstrate how structural bioinformatics, AI-based fold prediction, and traditional experiments complement each other and can shed light on the makeup of this important protein, both in SARS-CoV-2 and related viruses. Our method can be generalized for other multi-domain proteins, so we describe it in detail.

Our prediction-based approach reveals a previously undescribed folded domain, which we could confirm experimentally. Our research also suggests a potential function of the nidovirus-wide conserved domain Y1: This domain may be involved in the assembly of nsp3, nsp4, and nsp6 into the hexameric pore, which was discovered by electron tomography and exports RNA into the cytosol. The Y1-hexamer, however, could not be expressed and purified on its own. We also provide a revised domain segmentation and nomenclature of nsp3 domains based on a compilation of previous research and our own findings.

## Introduction and background

### Coronavirus biology

Coronavirus genomes encode several non-structural proteins (nsps), which permit the virus to deregulate the host immune system and to reproduce within the host cell [1]. Upon infection, the non-structural proteins are translated together as the large polyproteins pp1a and pp1ab, where the latter one is translated as a consequence of a ribosomal frame shift [2]. The polyprotein pp1a consists of 10 nsps, whereas pp1ab comprises 16 proteins. The polyproteins are cleaved into functional non-structural proteins by the viral papain-like proteases encoded in nsp3 and the main protease (nsp5) [3]. Depending on the species, nsp3 has one or two papain-like protease domains [4]. Without these proteases the infection cycle cannot progress, making nsp3 an important drug target in the fight against COVID-19. In this work, we focus on the murine hepatitis virus (MHV) and the two sarbecoviruses SARS-CoV-1 and SARS-CoV-2, which are of great interest due to the COVID-19 pandemic. All three belong to the genus *Betacoronavirus*, which is together with *Alphacoronavirus*, *Gammacoronavirus*, and *Deltacoronavirus* part of the subfamily *Orthocoronavirinae*. Those belong to the family of *Coronaviridae*, which in turn belongs to the order *Nidovirales* and finally to the positive single stranded RNA viruses.

Non-structural protein 3 (nsp3) accounts for 20% of the viral RNA, with 2006 amino acid residues in MHV, 1922 in SARS-CoV-1, and 1945 in SARS-CoV-2, respectively. Depending on the species, these residues form up to 16 domains (Fig. 1), including two transmembrane helices, which attach the protein to the membrane of double membrane vesicles (DMVs) [5] inside the host cell. These vesicles originate from endoplasmic reticulum during infection and house the replication of the viral genome [5]. While most regions of nsp3 are on the cytoplasmic side, one short region extends into the vesicle’s lumen (Figure 1 b) and interacts there with the lumenal domain of nsp4 in order to mediate double membrane vesicle formation [6]. Along with nsp4 and nsp6, nsp3 assembles into large complexes with a sixfold symmetry which function as pores, exporting the newly replicated viral RNA into the cytosol [7]. Subsequently, the RNA is packed into nucleocapsid proteins and is assembled with the other structural proteins into new virions, ready for the next infection [8].

**Figure 1.**
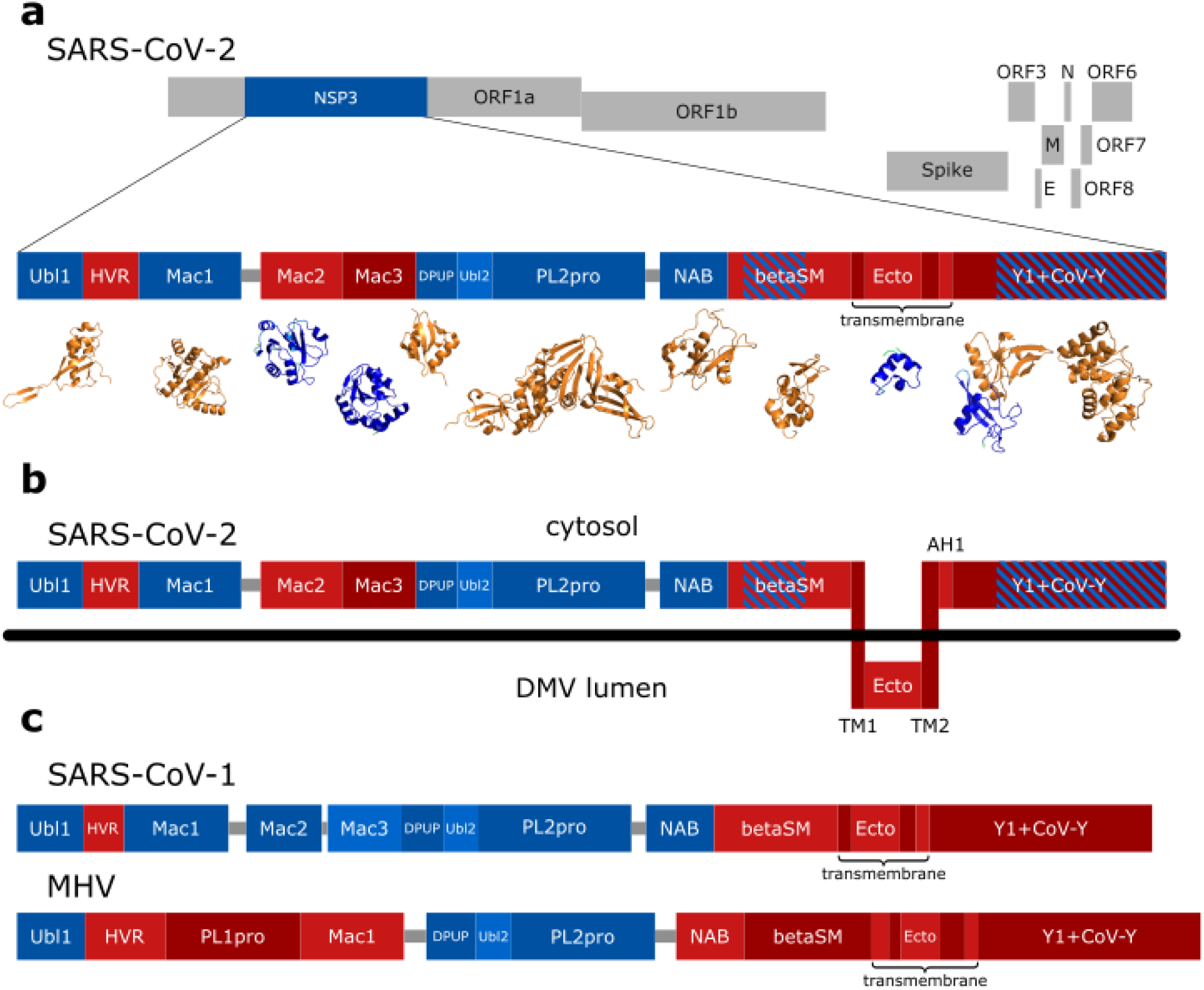
Domain overview of nsp3 from both sarbecoviruses and MHV. (a): Position and size of nsp3 on the polyprotein, as well as all domains (blue/red boxes) and larger linkers (grey lines). Blue domains are associated with experimentally solved structures in the PDB; red domains are not experimentally solved; red domains with blue stripes are solved partially. Depicted 3D structures in orange are from the PDB (see Table 1 for PDB codes), while the other structures are predicted by AlphaFold2 and coloured according to their pLDDT, with blue representing high confidence with pLDDT values greater than 90. (b): Membrane topology of SARS-CoV-2, based on the results of Oostra et al. [5]. (c): Domains of SARS-CoV-1 and MHV. Depicted domain ranges are mentioned in the results as preliminary ranges and are listed in Table S1 of the supplementary information. Domain ranges of SARS-CoV-1 are according to Lei et al. [4]; ranges of SARS-CoV-2 and MHV are based on sequence alignments with those of SARS-CoV-1 and experimental structures.

### Non-structural-protein 3 domains

The N-terminal domain of nsp3 is known as **ubiquitin like domain 1 (Ubl1)** and interacts with the nucleocapsid protein [9]. It is followed by a long, disordered domain connecting Ubl1 with the rest of nsp3, the **hypervariable region**. The next domain of nsp3 in MHV is the papain-like protease **PL^pro^**, performing the cleavage of nsp1 from the polyproteins, whereas in *Sarbecovirus* this domain is absent [4]. **Macrodomain 1 (Mac1)**, also known as ADP-ribose phosphatase, was shown to reverse the human PARP14-derived ADP-ribosylation [10], a mechanism involved in anti-viral defence, making Mac1 another well-researched nsp3 drug target [11]. Macrodomain 1 is followed by a linker and a region preceding PL2^pro^. For *Sarbecovirus* nsp3, this region was previously known as the SARS-unique domain, consisting of **macrodomain 2 (Mac2)**, **macrodomain 3 (Mac3)** and the **domain preceding Ubl2 and PL2^pro^ (DPUP)**. However, only Mac2 and Mac3 are unique to *Sarbecovirus*, while MHV possess a DPUP-like region [12]. Next comes the **ubiquitin-like domain 2 (Ubl2)**, which is seen as a subdomain of **PL2^pro^**. This (in MHV second) papain-like protease is an essential enzymatic domain and therefore a drug target [4,13]. It cleaves the polyproteins between nsp2 and nsp3, between nsp3 and nsp4, and in *Sarbecovirus* additionally between nsp1 and nsp2 [4]. PL2^pro^ is separated from the consecutive **nucleic acid binding domain (NAB)** by a ∼30 residue linker. However, as we will show below, this “linker” is a folded and conserved domain specific to *Betacoronavirus*. The last third of nsp3 consists of a **betacoronavirus-specific marker domain (βSM)**, the transmembrane region with a lumenal domain (known as **ectodomain**), and the C-terminal region [4]. This C-terminal region consists of the **nidovirus-conserved domain of unknown function (Y1)** and **CoV-Y** [14], whereas two published structures (PDB codes 7RQG [15]; 8F2E [16]) and a recent paper [16] suggest to divide both domains into two subdomains.

Currently, experimentally determined structures are only available for the domains Ubl1, Mac1, Mac2, Mac3, DPUP, Ubl2, PL2^pro^, NAB, part of βSM, the second half of Y1, and CoV-Y (see Fig. 1). The majority of the available PDB depositions features either Mac1 or PL2^pro^, as pointed out by the Coronavirus Structural Task Force [17,18]. The atomic structures of the transmembrane region and part of the C-terminal region were not experimentally elucidated. However, novel structure prediction methods enable a first glimpse into their potential folds.

### AI-based fold prediction

In August 2021, the structure prediction software AlphaFold2 [19] became publicly available and enabled a first look into the 3D structure of the undetermined nsp3 domains. AlphaFold2 is based on convolutional neural networks with a two-tower transformer architecture, utilizing co-evolution of sequentially remote but spatially close amino acids. The networks were trained on the PDB and multi-sequence alignments of protein sequences [20]. AlphaFold2 predictions closely resemble experimentally determined structures in terms of backbone fold, especially in the case of small and stably folded proteins [21,22] and come with a confidence score for each residue. Although not intended, this confidence metric, known as the predicted local distance difference test (pLDDT), can be an indicator of disorder [23–25]. Another metric, the predicted alignment error (pAE), assists in the recognition and evaluation of subdomains and their relative alignment [26].

In our work, we utilized AlphaFold2 to predict the structures of all regions of nsp3 from MHV, SARS-CoV-1 and SARS-CoV-2, which were used along experimentally determined structures and the provided metrics from AlphaFold2 to determine domain ranges. This led to the discovery of a new domain and the hexameric assembly of the nsp3 C-terminus, followed by experimental validation. In the following, we will demonstrate (A) how we used AlphaFold2 iteratively for construct design and for definition of domain ranges; (B) investigation and validation of the linker domain; and finally (C) analysis of the predicted Y1+CoV-Y hexamer.

## Results

Non-structural protein 3 (nsp3) is a large multidomain protein present in all coronaviruses. Together with other proteins it assembles into a hexameric complex that exports new copies of the viral genome into the cytosol and plays therefore an essential role in the viral replication cycle.

We unified the domain boundaries of the individual nsp3 domains, as these were based on few experimental data and were contradictory between publications. During this process, we developed a method for the detection of domains in a multi-domain protein based on features of AlphaFold2. With this, we discovered an undescribed domain and validated its fold via a SAXS (small-angle X-ray scattering) experiment. Investigation of the C-terminal Y1 domain led to the hypothesis of Y1 being a major driving force in the assembly of the hexameric RNA-exporting channel. This hypothesis is supported by our bioinformatical analysis, but requires further experimental assessment.

### Utilizing AlphaFold2 for domain boundary determination and construct design

#### Preliminary domain ranges

The number of domains and the exact domain boundaries for nsp3 of SARS-CoV-1 and the related murine hepatitis virus (MHV) changed a lot over the time [14]. This led to contradictory information in the current literature and to partially incomplete gene annotations, especially in the case of SARS-CoV-2. Since the definition of the last domain ranges [4], numerous experimentally determined structures of nsp3 domains emerged, which enable a more complete and accurate domain definition. Furthermore, structure prediction gives insight into currently unresolved regions of nsp3. Here, we aimed at assigning domain ranges to all nsp3 domains of three distinct viruses and unified existing naming conventions. To accomplish this, we utilized recent experimental structures as well as AI-based fold prediction.

The available experimental structures cover only a fraction of nsp3. In order to augment domain ranges via structure prediction and investigate unresolved domains, input sequences for AlphaFold2 [19] had to be designed. These input sequences were based on sequences from experimentally determined structures and on the domain ranges listed in Lei et al. [4] for SARS-CoV-1 and MHV, as well as on the genome annotation entry YP_009742610.1 from NCBI for SARS-CoV-2. Additional information was provided by transmembrane domain predictions via TMHMM 2.0 [27]. Together, this was sufficient to define preliminary domain ranges, which cover all ∼2000 residues of nsp3.

Since the number of experimentally solved domains was limited for SARS-CoV-2 and MHV, we first defined the preliminary domain ranges for SARS-CoV-1. The list was then completed for the other two viruses via sequence alignments against each SARS-CoV-1 domain (see Table S1 of the supplementary information for all preliminary domain ranges).

#### Sequence alignments of preliminary domain ranges

In order to estimate the similarity of domains between both sarbecoviruses and MHV, sequences of the preliminary ranges were compared in local pairwise sequence alignments (see Table S2 of the supplementary information). Sequence identities were mostly high between the two sarbecoviruses and lower between sarbecoviruses and MHV. From the 18 domains and large linkers, four regions are assumed to be intrinsically disordered [4]. From those, however, only three (linker Mac1-Mac2, hypervariable region, and betacoronavirus-specific marker domain) show relatively low identities (41% to 69% between sarbecoviruses). With 80%, the sequence identity of the linker between PL2^pro^ and NAB is comparable to the ordered regions with identities of 71% to 90%. The two C-terminal domains Y1 and CoV-Y are among the domains with the highest identity (88.1% for Y1 and 90.2% for CoV-Y between sarbecoviruses), which are studied together with the PL2^pro^- NAB linker in the following sections.

#### Comparison between experimentally known structures and AlphaFold2 predictions

The preliminary domain ranges (see Table S1 of the supplementary information) were used to generate input sequences for AlphaFold2 [19], covering the entire nsp3 residue range. If an experimentally determined structure of a domain was available, the AlphaFold2 prediction was compared to it (Table 1 lists the respective root mean square deviations (RMSD)).

**Table 1.** Comparison of AlphaFold2 predictions to corresponding experimental structures with RMSD values. We used the rank 1 predictions from AlphaFold2 and calculated the RMSD with PyMOL [30]. In case of Mac1 and PL2^pro^ for SARS-CoV-2, where a great number of published structures exists, we compared to the structures with the highest resolution and picked the one with the lowest RMSD. For all other domains, we tested all PBD entries and listed here the lowest RMSD, with the tested PDB structure behind the RMSD value. In case of 8F2E, atoms were removed prior alignment to match the exact subdomains.

| Name | Abbreviation | SARS-CoV-2 | SARS-CoV-1 | MHV |
| --- | --- | --- | --- | --- |
| Ubiquitin-like domain 1 | Ubl1 | 0.5 Å (7KAG) | 1.3 Å (2GRI) | 2.7 Å (2M0A) |
| Macrodomain 1 | Mac1 | 0.3 Å (7KQP) | 0.3 Å (2ACF) | - |
| Macrodomain 2 | Mac2 | - | 0.4 Å (6YXJ) | - |
| Macrodomain 3 | Mac3 | - | 0.7 Å (2JZD) | - |
| Domain preceding Ubl2 and PL2<sup>pro</sup> | DPUP | 0.3 Å (7THH) | 0.6 Å (2KAF) | 0.4 Å (4YPT) |
| Ubiquitin-like domain 2 | Ubl2 | 0.2 Å (7D6H) | 0.2 Å (2FE8) | 0.3 Å (5WFI) |
| Papain-like protease 2 | PL2<sup>pro</sup> | 0.7 Å (7D6H) | 0.5 Å (5TL7) | 0.5 Å (5WFI) |
| Nucleic-acidic-binding domain | NAB | 0.4 Å (7LGO) | 0.8 Å (2K87) | - |
| Betacoronavirus-specific marker domain | βSM | 0.9 Å (7T9W) | - | - |
| Nidovirus-conserved domain of unknown function subdomain b | Y1b | 0.4 Å (8F2E) | - | - |
| Coronavirus-specific C-terminal domain | CoV-Ya | 0.4 Å (8F2E) | - | - |
| Coronavirus-specific C-terminal domain | CoV-Yb | 0.4 Å (7RQG) | - | - |

Most cases show RMSD values below 1 Å and high per-residue confidence (high predicted local distance difference test (pLDDT) values) across the whole structure. The two exceptions, Ubl1 domains from SARS-CoV-1 and MHV, contain a flexible N-terminus [28,29], which had pLDDT values below 50. Although small misalignments were observed for loops and secondary structure elements, the number and presence of such elements was correct in all cases listed in Table 1.

#### Classification into regions of order and disorder

AlphaFold2 predictions come with a confidence score (pLDDT; predicted local distance difference test) and a predicted alignment error (pAE) for each residue, which evaluate the local and global folding, respectively. We combined pLDDT and the predicted 3D fold to differentiate between regions of order and disorder. Individual domains within one large segment of order are distinguished via the pAE. Ordered folds are characterised by multiple secondary structure elements and pLDDT values above 80. Loops that are either surrounded by such regions or show high pLDDT values are also considered ordered. Potentially disordered regions on the other hand are lacking secondary structure elements and have pLDDT values below 50. Figure 2 shows our decision tree based on these criteria, used to classify each region into potentially ordered or disordered.

**Figure 2.**
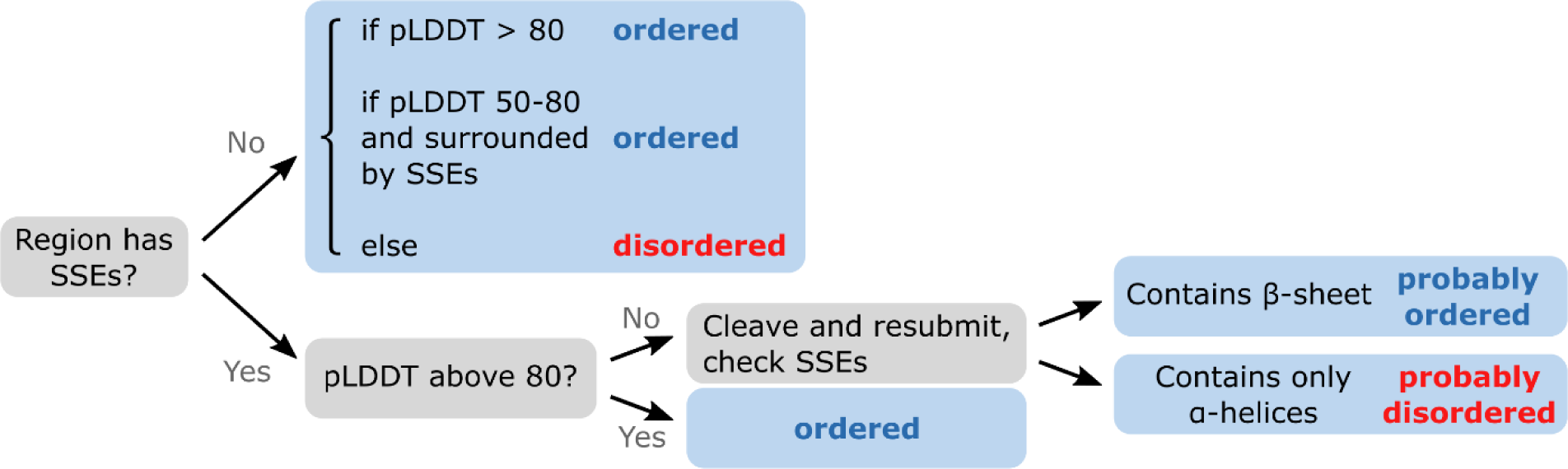
Decision tree for the classification of regions from fold-predictions into ordered or disordered. SSEs stands for “secondary structure elements”; pLDDT for “predicted local distance difference test”. Multiple iterations are only required if SSEs with low pLDDT or large low pLDDT loops at the termini are present, where cleavage must take place.

Another property for differentiation is the orientation of the termini. Aligning multiple fold-predictions leads to a near perfect overlap of ordered sections, while disordered termini point away from the fold in random orientations. Figure 3a illustrates this phenomenon well on predictions of the domain Ubl1 with its disordered N-terminus. The predicted, stretched-out “barbed wire” conformations are furthermore inconsistent with normal main-chain torsions, as described by Williams et al. [25].

**Figure 3.**
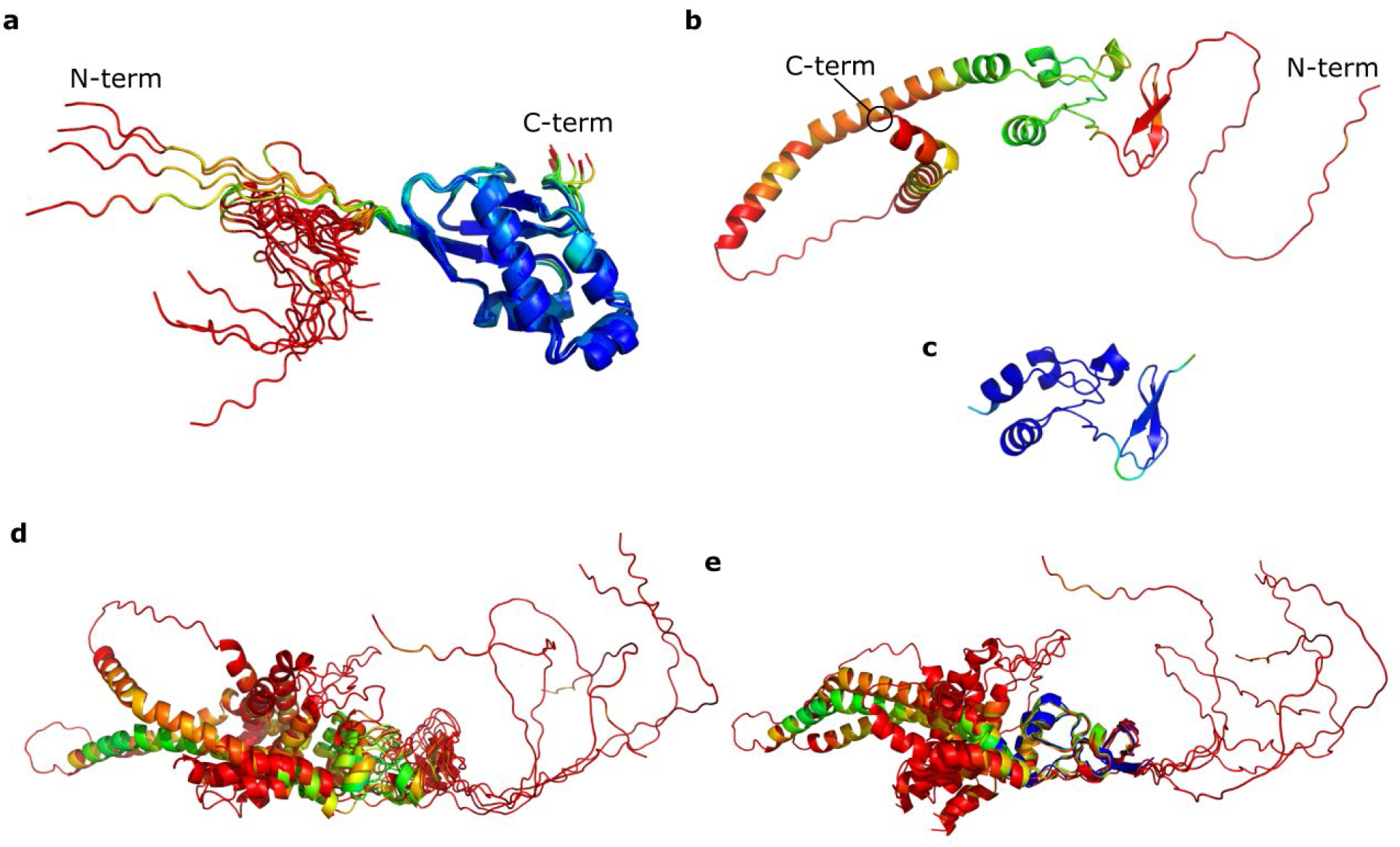
Ordered and disordered region in relation to confidence scores in predicted models. Residues are colored according to their pLDDT, with blue representing high confidence and values above 90, red representing low confidence with values below 50, and other colors representing values inbetween. (a): Ensemble of twenty predicted models emphasizing the difference between disorder (red, left) and order (blue, right). Predictions of the Ubl1 domain from SARS-CoV-1 via AlphaFold2 by running five models for a total of four seeds. While the secondary structure elements overlap clearly, the disordered N-terminus to the left varies drastically between models. (b): Prediction of complete βSM domain from SARS-CoV-2 with low confidence secondary structure elements and large disordered termini. (c): Prediction of a cropped βSM domain sequence lacking the termini, showing an increased overall pLDDT. (d): Ensemble of five predicted models aligned to the rank1 model, with the central fold recognizable. (e): Same ensemble as in (d) aligned to the experimentally determined structure 7T9W colored in blue. Disordered regions, despite being predicted as helices, point away in various directions.

A last special case covers ordered regions which are predicted with well-defined secondary structure elements but with low pLDDT. The prediction of the βSM domain is a good example (Figure 3b). The input sequence covers a large disordered N-terminus, an ordered central part, and a large disordered C-terminus. A low pLDDT is assigned to the termini, thus correctly implying intrinsic disorder for that region. However, these termini influence the confidence of the central fold negatively. Cropping these termini and resubmitting the sequence of just the ordered part leads to a high confidence prediction (Figure 3c). Furthermore, the new prediction is more accurate with the RMSD to the experimentally solved structure (PDB: 7T9W) dropping from 0.9 Å to 0.5 Å. The C-terminal helices keep low pLDDT values even after sequence cropping, which classifies it as potentially disordered.

#### Final determination of domain boundaries

The iterative method of predicting, classifying, cropping, and resubmitting for prediction was utilized to identify all regions of order and disorder within nsp3. The boundaries between ordered and disordered regions set the final domain ranges and in case of large segments of order, the predicted alignment error (pAE) was utilized for domain separation. The resulting pAE matrices are contained in Figure S1 of the supplementary information.

One large region of order comprised the domains Mac3, DPUP, Ubl2, PL2^pro^, a newly discovered domain, and NAB. Because is was not possible to distinguish between Ubl2, PL2^pro^, and the new fold following PL2^pro^ (from here named “linker domain”) in the pAE matrix, ranges from literature and experimentally determined structures were used to define domain boundaries. The newly discovered linker domain is further examined in section B.

The final domain ranges are listed in Table 2. Noteworthy discoveries are a folded helix in the otherwise disordered hyper variable region for MHV (nsp3 residues 230-241); the aforementioned linker domain between PL2^pro^ and NAB; a fold in the ectodomain; and a full prediction of Y1. None of these have been experimentally observed before. All regions of potential disorder in nsp3 make up 357 residues (18.4%) in SARS-CoV-2, 344 (17.9%) in SARS-CoV-1, and 506 (25.2%) in MHV.

**Table 2.**
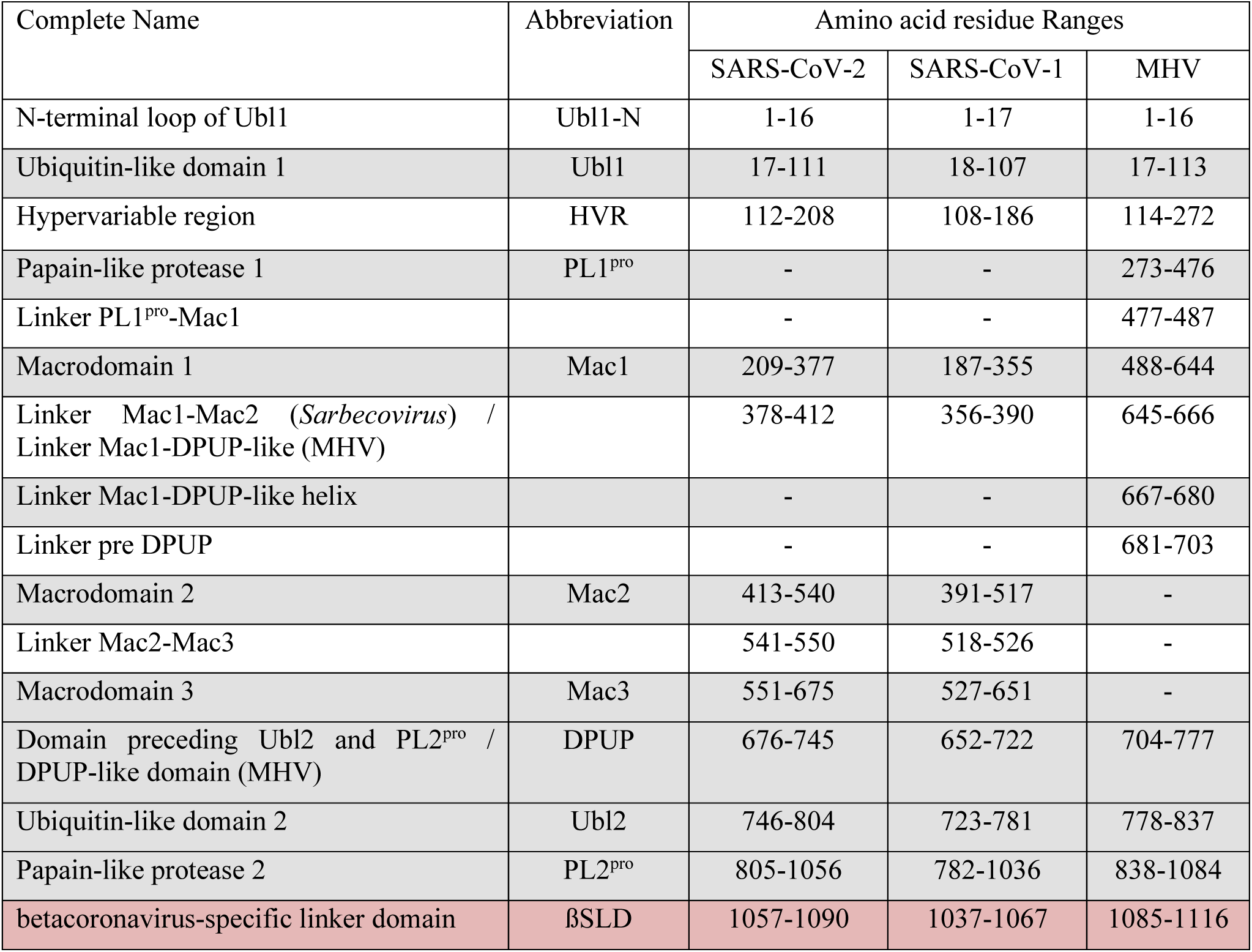
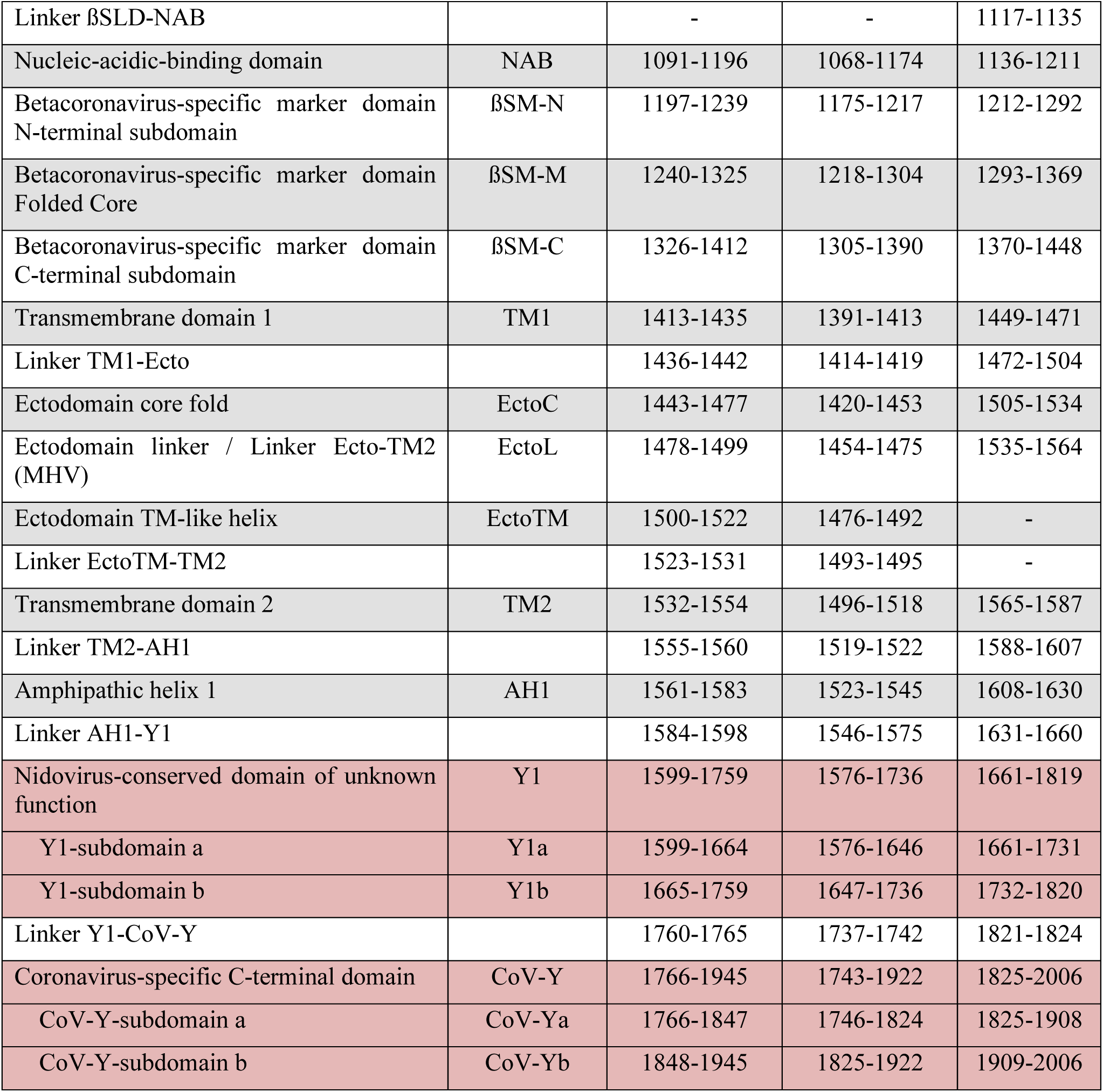
Ranges of all domains and linkers determined by combining AlphaFold2 and experimentally solved structures. Shaded entries are predicted to have a defined structure, while non-shaded entries describe predicted regions of disorder. Domains discussed in more detail are shaded in red. Note that these domain boundaries are based on predicted order and are useful for crystallization trials. For full biological functionality, some domains may require their surrounding linkers.

| Complete Name | Abbreviation | Amino acid residue Ranges |  |  |
| --- | --- | --- | --- | --- |
|  |  | SARS-CoV-2 | SARS-CoV-1 | MHV |
| N-terminal loop of Ubl1 | Ubl1-N | 1-16 | 1-17 | 1-16 |
| Ubiquitin-like domain 1 | Ubl1 | 17-111 | 18-107 | 17-113 |
| Hypervariable region | HVR | 112-208 | 108-186 | 114-272 |
| Papain-like protease 1 | PL1<sup>pro</sup> | - | - | 273-476 |
| Linker PL1<sup>pro</sup>-Mac1 |  | - | - | 477-487 |
| Macrodomain 1 | Mac1 | 209-377 | 187-355 | 488-644 |
| Linker Mac1-Mac2 (*Sarbecovirus*) /<br>Linker Mac1-DPUP-like (MHV) |  | 378-412 | 356-390 | 645-666 |
| Linker Mac1-DPUP-like helix |  | - | - | 667-680 |
| Linker pre DPUP |  | - | - | 681-703 |
| Macrodomain 2 | Mac2 | 413-540 | 391-517 | - |
| Linker Mac2-Mac3 |  | 541-550 | 518-526 | - |
| Macrodomain 3 | Mac3 | 551-675 | 527-651 | - |
| Domain preceding Ubl2 and PL2<sup>pro</sup> /<br>DPUP-like domain (MHV) | DPUP | 676-745 | 652-722 | 704-777 |
| Ubiquitin-like domain 2 | Ubl2 | 746-804 | 723-781 | 778-837 |
| Papain-like protease 2 | PL2<sup>pro</sup> | 805-1056 | 782-1036 | 838-1084 |
| betacoronavirus-specific linker domain | βSLD | 1057-1090 | 1037-1067 | 1085-1116 |
| Linker βSLD-NAB | | - | - | 1117-1135 |
| Nucleic-acidic-binding domain | NAB | 1091-1196 | 1068-1174 | 1136-1211 |
| Betacoronavirus-specific marker domain N-terminal subdomain | βSM-N | 1197-1239 | 1175-1217 | 1212-1292 |
| Betacoronavirus-specific marker domain Folded Core | βSM-M | 1240-1325 | 1218-1304 | 1293-1369 |
| Betacoronavirus-specific marker domain C-terminal subdomain | βSM-C | 1326-1412 | 1305-1390 | 1370-1448 |
| Transmembrane domain 1 | TM1 | 1413-1435 | 1391-1413 | 1449-1471 |
| Linker TM1-Ecto |  | 1436-1442 | 1414-1419 | 1472-1504 |
| Ectodomain core fold | EctoC | 1443-1477 | 1420-1453 | 1505-1534 |
| Ectodomain linker / Linker Ecto-TM2 (MHV) | EctoL | 1478-1499 | 1454-1475 | 1535-1564 |
| Ectodomain TM-like helix | EctoTM | 1500-1522 | 1476-1492 | - |
| Linker EctoTM-TM2 |  | 1523-1531 | 1493-1495 | - |
| Transmembrane domain 2 | TM2 | 1532-1554 | 1496-1518 | 1565-1587 |
| Linker TM2-AH1 |  | 1555-1560 | 1519-1522 | 1588-1607 |
| Amphipathic helix 1 | AH1 | 1561-1583 | 1523-1545 | 1608-1630 |
| Linker AH1-Y1 |  | 1584-1598 | 1546-1575 | 1631-1660 |
| Nidovirus-conserved domain of unknown function | Y1 | 1599-1759 | 1576-1736 | 1661-1819 |
| Y1-subdomain a | Y1a | 1599-1664 | 1576-1646 | 1661-1731 |
| Y1-subdomain b | Y1b | 1665-1759 | 1647-1736 | 1732-1820 |
| Linker Y1-CoV-Y |  | 1760-1765 | 1737-1742 | 1821-1824 |
| Coronavirus-specific C-terminal domain | CoV-Y | 1766-1945 | 1743-1922 | 1825-2006 |
| CoV-Y-subdomain a | CoV-Ya | 1766-1847 | 1746-1824 | 1825-1908 |
| CoV-Y-subdomain b | CoV-Yb | 1848-1945 | 1825-1922 | 1909-2006 |

During the classification into ordered and potentially disordered regions, few domains were divided into new subdomains and linkers, which extends the number of ranges compared to previous literature. One such case is the division of ßSM into ßSM-N (N-terminal linker), ßSM-M (ordered fold), and ßSM-C (C-terminal linker). While the terminal linkers are likely disordered due to low pLDDT and low sequence identity in alignments, ßSM-M resembles the experimentally determined structure 7T9W. As we show in section B, the domain between PL2^pro^ and NAB is betacoronavirus specific, hence the name “betacoronavirus-specific linker domain”.

The final domain ranges of all three viruses were compared for sequence similarity and additionally for similarity between predicted structures by calculating their RMSD values (Table 3). The predictions for both sarbecoviruses show high similarity with all RMSD values below 0.8 Å (excluding the transmembrane domains). Comparing both viruses to MHV, however, only seven out of eleven domains (excluding disordered and transmembrane domains) show high similarity with RMSDs of 1.1 Å or less. With RMSD values above 7 Å, the prediction of MHV’s ectodomain stands out, which is explored in section C.

**Table 3.**
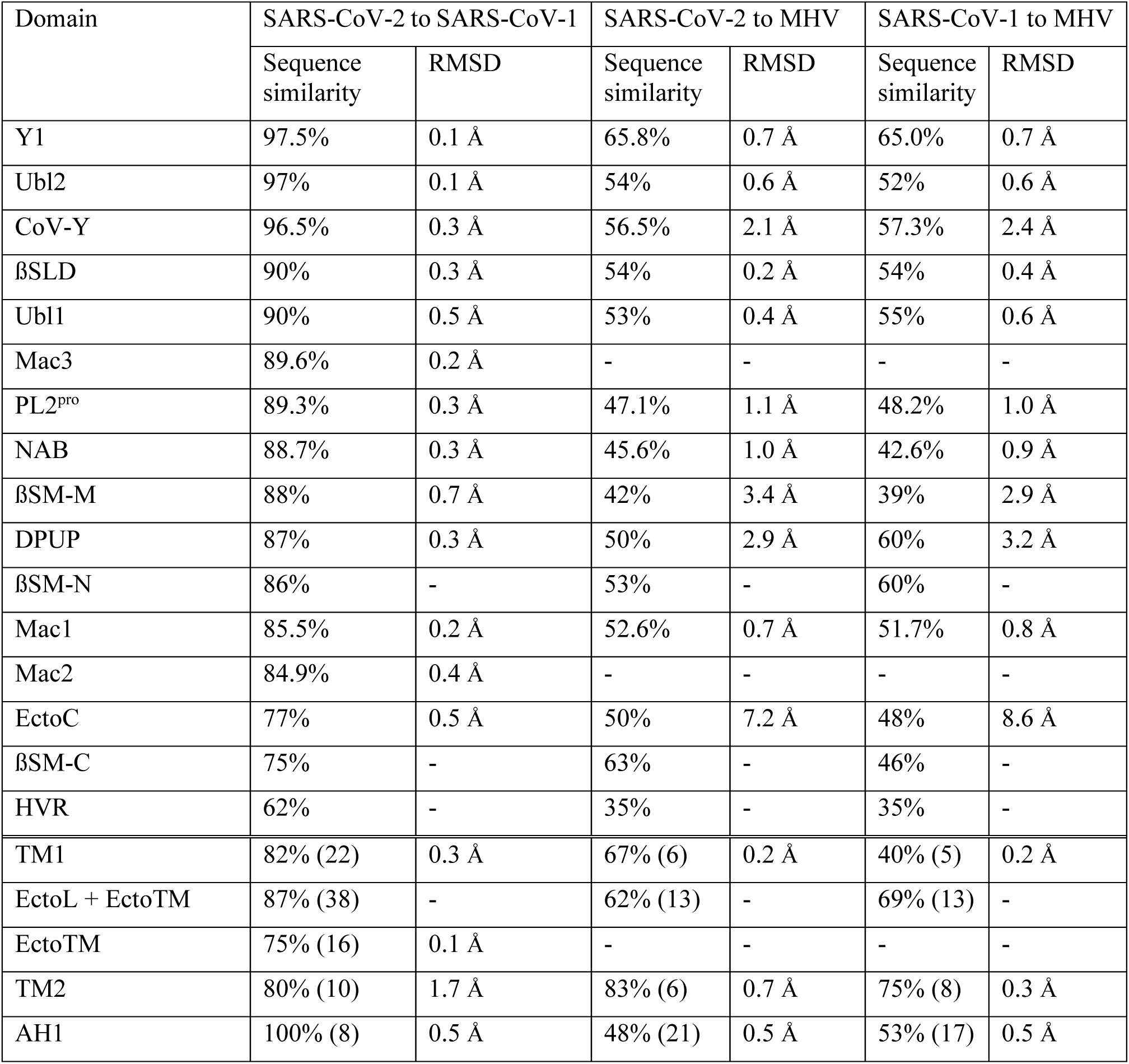
Sequence similarities and RMSD values between pairs of MHV, SARS-CoV-1 and SARS-CoV-2. Only domains predicted to fold into a defined structure and large regions of disorder are listed. Short linkers are omitted. RMSD values are calculated for defined folds with PyMOL [30]. Results are sorted by decreasing sequence similarity between both sarbecoviruses. Domains of the transmembrane region are listed below and are not sorted, since only short alignments were found. For these cases, the alignment-length is given in parentheses.

| Domain | SARS-CoV-2 to SARS-CoV-1 |  | SARS-CoV-2 to MHV |  | SARS-CoV-1 to MHV |  |
| --- | --- | --- | --- | --- | --- | --- |
|  | Sequence similarity | RMSD | Sequence similarity | RMSD | Sequence similarity | RMSD |
| Y1 | 97.5% | 0.1 Å | 65.8% | 0.7 Å | 65.0% | 0.7 Å |
| Ubl2 | 97% | 0.1 Å | 54% | 0.6 Å | 52% | 0.6 Å |
| CoV-Y | 96.5% | 0.3 Å | 56.5% | 2.1 Å | 57.3% | 2.4 Å |
| βSLD | 90% | 0.3 Å | 54% | 0.2 Å | 54% | 0.4 Å |
| Ubl1 | 90% | 0.5 Å | 53% | 0.4 Å | 55% | 0.6 Å |
| Mac3 | 89.6% | 0.2 Å | - | - | - | - |
| PL2<sup>pro</sup> | 89.3% | 0.3 Å | 47.1% | 1.1 Å | 48.2% | 1.0 Å |
| NAB | 88.7% | 0.3 Å | 45.6% | 1.0 Å | 42.6% | 0.9 Å |
| βSM-M | 88% | 0.7 Å | 42% | 3.4 Å | 39% | 2.9 Å |
| DPUP | 87% | 0.3 Å | 50% | 2.9 Å | 60% | 3.2 Å |
| βSM-N | 86% | - | 53% | - | 60% | - |
| Mac1 | 85.5% | 0.2 Å | 52.6% | 0.7 Å | 51.7% | 0.8 Å |
| Mac2 | 84.9% | 0.4 Å | - | - | - | - |
| EctoC | 77% | 0.5 Å | 50% | 7.2 Å | 48% | 8.6 Å |
| βSM-C | 75% | - | 63% | - | 46% | - |
| HVR | 62% | - | 35% | - | 35% | - |
| TM1 | 82% (22) | 0.3 Å | 67% (6) | 0.2 Å | 40% (5) | 0.2 Å |
| EctoL + EctoTM | 87% (38) | - | 62% (13) | - | 69% (13) | - |
| EctoTM | 75% (16) | 0.1 Å | - | - | - | - |
| TM2 | 80% (10) | 1.7 Å | 83% (6) | 0.7 Å | 75% (8) | 0.3 Å |
| AH1 | 100% (8) | 0.5 Å | 48% (21) | 0.5 Å | 53% (17) | 0.5 Å |

In contrast to high structural similarity between MHV and both sarbecoviruses, the sequence similarities are relatively low with values from 42.6% to 65.8%. The C-terminal domains Y1 and CoV-Y stand out with high sequence similarities of up to 97.5% despite containing more than 150 residues, indicating an important function. Surprisingly, the linker domain is listed at position four with higher similarity than PL2^pro^ and the macrodomains. Its structure and conservation among coronaviruses are explored in the next section.

### Experimental validation of the Betacoronavirus-specific linker domain (βSLD)

#### Structure prediction of linker domain between PL2^pro^ and NAB domain

The linker between PL2^pro^ and NAB stands out due to its high sequence similarity of 90% between SARS-CoV-2 and SARS-CoV-1, higher than the similarities of most other folded domains (Table 3). Additionally, AlphaFold2 predicts a stable fold consisting of 34 residues for SARS-CoV-2, forming two pairs of antiparallel β-sheets connected by an 8-residue loop (Figure 4). This loop was predicted to form a helix-like structure, which could potentially form due to a restriction in its conformational space induced by Pro1076 and two hydrogen bonds within its backbone. A similar helix-like structure is found at the N-terminus, also containing a proline and two internal hydrogen bonds. The fourth β-sheet in the sequence Nterm-β1-β2-loop-β3-β4-Cterm is connected via two hydrogen bonds to β1 in parallel direction, leading to a compact fold. The sheet β1 is connected to β2 via hydrogen bonds in the backbone and additionally via the Thr1063-Thr1072 and Tyr1064-Glu1073 hydrogen bonds (Figure 4). Ramachandran analysis via MolProbity [31] shows only Thr1058 as an outlier, while 26 out of 32 residues were in favoured regions (Ramachadran plots in Figure S2 of the supplementary information).

**Figure 4.**
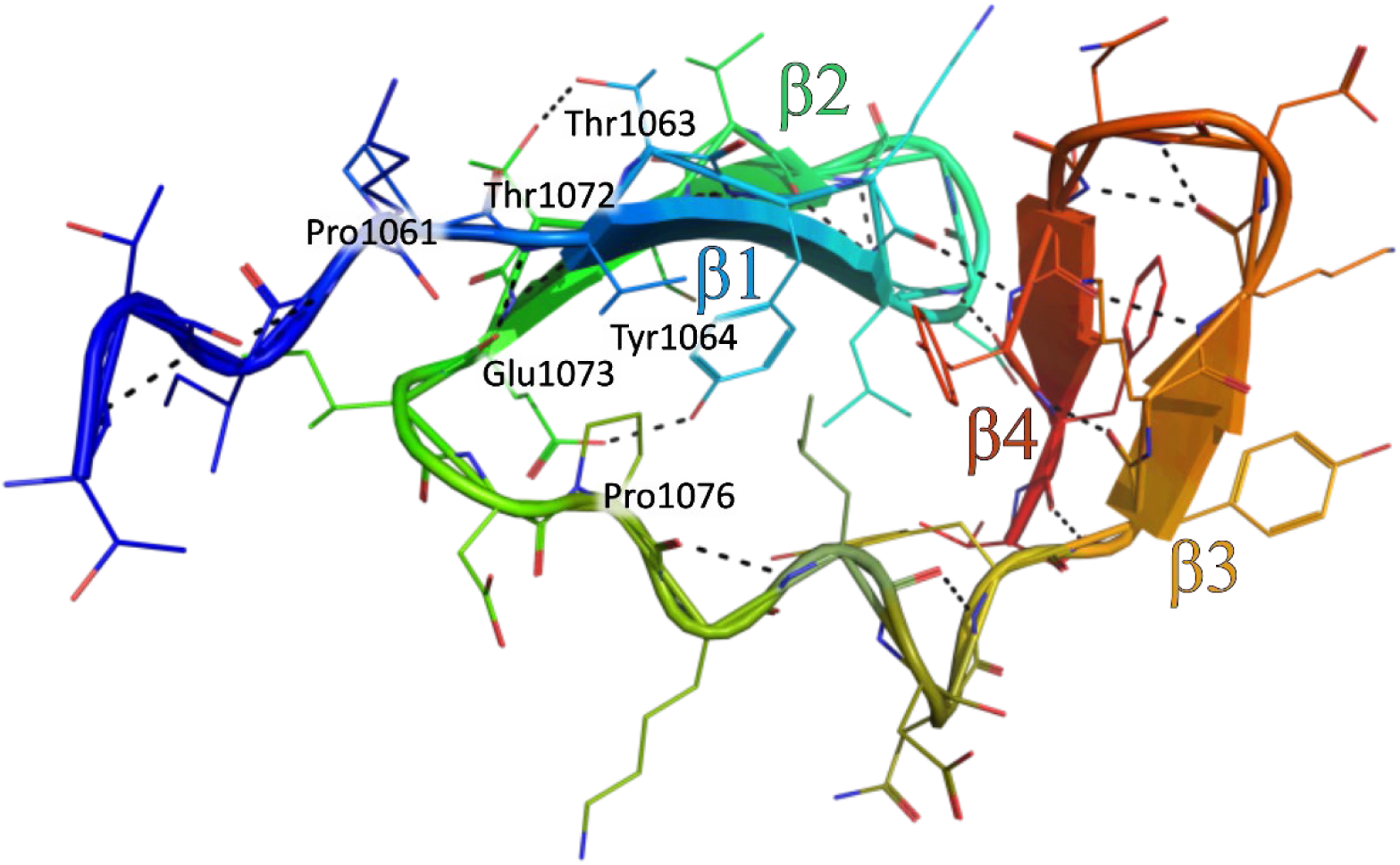
AlphaFold2 structure prediction of the SARS-CoV-2 nsp3 linker domain. N-terminus is coloured in blue, C-terminus in red. Hydrogen bonds are depicted as black dotted lines. The labelled prolines potentially restrict the conformational freedom of their loops, with Pro1076 being located in the central loop. The other labelled residues interact via hydrogen bonds with each other. Residue numbers are based on SARS-CoV-2 nsp3 sequence.

#### Conservation of the linker domain

The sequence of this linker domain from MHV has a high sequence similarity to the linker domain from both sarbecoviruses (54%). A BLAST search yielded additional hits within *Betacoronavirus*, but no hits outside of this genus.

Local pairwise and multiple sequence alignments between the sequence from SARS-CoV-2 and sequences from 16 betacoronaviruses revealed conserved residues (Figure 5), namely Leu1066, Asp1067, Pro1076, Tyr1088, and Thr1090 (residue numbers based on SARS-CoV-2 nsp3). Furthermore, the residues 1064, 1081 and 1089 are always Phe or Tyr; Leu1078 is also mostly conserved. From 34 residues, 14 retain similar chemical properties, including fully conserved ones (Figure 5). From those, 12 are in a loop or at the edge of a β-sheet right next to a loop, indicating a high selection pressure on the loop regions. The structure predictions show in all cases a highly similar fold with RMSD values below 0.2 Å.

**Figure 5.**
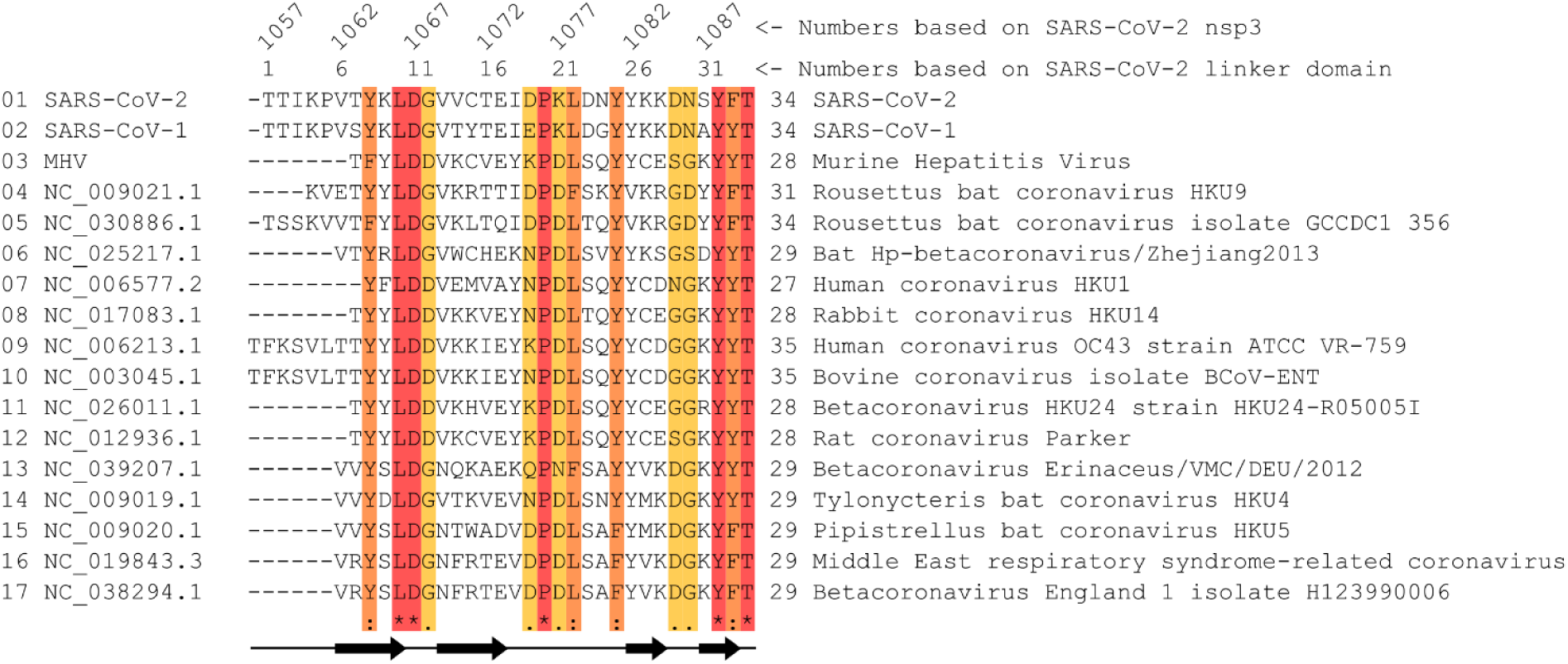
Multiple sequence alignment between the linker domain of various betacoronaviruses. The sequences for the linker domains were first identified with a local pairwise sequence alignment between the linker domain sequence of SARS-CoV-2 and orf1ab sequence of the respective virus. Afterwards, the identified sequences were used in a global multiple sequence alignment, resulting in this figure. Residues conserved in all examined viruses are shown in red, residues with strongly similar chemical properties at the same position are highlighted in orange, and weakly similar properties are shown in yellow. The secondary structure elements are indicated at the bottom.

Additional sequence alignments with the unclassified shrew coronavirus, alpha- (24 viruses), gamma- (5 viruses), and deltacoronaviruses (10 viruses) revealed no hits within the region of nsp3. However, several hits were observed in the region of endoRNAse, known as nsp15 in SARS-CoV-2. Structure predictions reveal a similar fold as for the linker domain, but with an additional helix before the third β-sheet. Details are in the supplementary information section B. Since no sequence homologues were identified outside of *Betacoronavirus*, we named this domain betacoronavirus-specific linker domain.

#### Experimental validation

To assess if the betacoronavirus-specific linker domain (βSLD) is in fact folded, single-crystal X-ray diffraction and small-angle X-ray scattering (SAXS) experiments were conducted. Due to the small size of the linker domain and the high confidence prediction of a close arrangement between this domain and PL2^pro^, a multidomain construct was designed. The construct comprised SARS-CoV-2 residues Thr1057 to Thr1090, including the domains Ubl2, PL2^pro^, and βSLD. Furthermore, the PL2^pro^ C111S mutant was used for better crystallization chances [13].

Crystallization trials led to thin crystals, from which data could be collected and processed into an electron density map. Unfortunately, the model building revealed that cleavage took place and only half of the construct assembled into the crystal. It was the undesired half covering Ubl2 and approximately half of PL2^pro^. Despite the surprising result that half of PL2^pro^ crystallizes, no useful information about the structure of βSLD could be gathered. It is unclear whether PL2^pro^ was involved in the proteolysis or other factors introduced a systematic cleavage. However, it is unlikely that PL2^pro^ is able to self-cleave *in vivo* due to its sequence-specificity [4] and the assembly of nsp3 into large hexameric complexes [7], as well as no reports about such behaviour. Analysis of the construct prior crystallization showed a band at the expected size. Also, remaining protein sample was used in a SAXS experiment, where cleavage of the construct could be excluded.

The SAXS result shows a good agreement between the relaxed AlphaFold2 prediction and the experimental data, with a *χ*2 of 0.98 (Fig. 6a) (The relaxed AlphaFold2 prediction is the most fitted structure found by the end of a single SREFLEX run [32]). The dimensionless Kratky plot suggests that the solution structure of Ubl2-PL2^pro^-βSLD is a rather compacted multidomain entity (Fig. 6b). It is highly unlikely that a long disordered tail exists, according to the flat plateau at the high scattering vector in the Kratky plot; instead, short flexible regions between domains are expected. The relaxed AlphaFold2 prediction also fits well against the envelop of the *ab initio* SAXS model (Fig. 6c).

**Figure 6.**
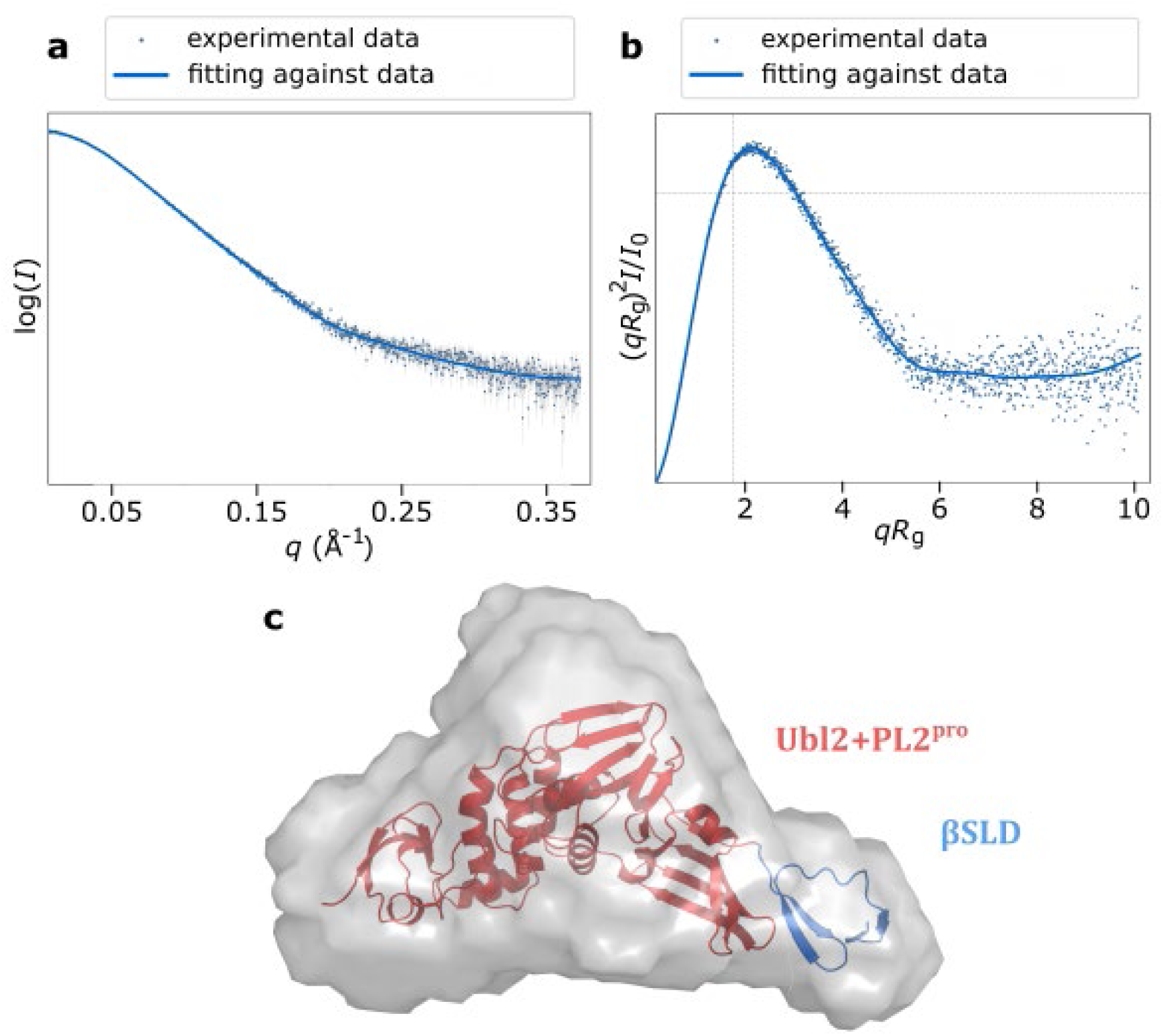
(a) The fitting result of relaxed AlphaFold2 model against the experimental SAXS data. Dots are the SAXS data with the relative errors. Solid line is the estimated scattering curve of the relaxed AlphaFold2 prediction. (b) The dimensionless Kratky plot. The peak maximum largely shifts away from the theoretical value for a compacted globular (marked by the two dashed grey lines) suggests the solution structure of Ubl2-PL2^pro^-βSLD is a multidomain protein. However, the convergence of the Kratky plot at higher q means there are no long flexible linkers between domains or at the N, C-termini of the entity. (c) A projection of the envelope of the ab initio model (grey volume) and the relaxed AlphaFold2 prediction (cartoon), with the linker domain at the right side in blue and the remaining part being Ubl2+PL2pro in red.

#### Nucleic acid binding domain and Betacoronavirus-specific marker domain

The subsequent domains from the betacoronavirus-specific linker domain, NAB (nucleic acid binding domain) and βSM (betacoronavirus-specific marker domain) are also betacoronavirus-specific [4,14]. However, a region similar to βSM was identified in gammacoronaviruses, known as γSM (gammacoronavirus-specific marker domain) [4]. Due to the large, disordered termini of βSM, we introduced a more specific nomenclature, with βSM-N (N-terminal link), βSM-M (folded central unit), and βSM-C (C-terminal link) with ranges in Table 2.

Sequence alignments within *Betacoronavirus* and structure predictions show that both, NAB and βSM-M, are present in all betacoronaviruses. Alignments between sequences from SARS-CoV-2 and 15 betacoronaviruses, excluding SARS-CoV-1, show sequence identities from 27% to 38% for NAB and 18% to 33% for βSM-M. NAB and βSM-M show also a high sequence similarity of 56% in an alignment of 25 residues, which could serve as a hint on their evolution.

In *Gammacoronavirus*, alignments with the Canada goose coronavirus showed the highest sequence identities, with 22% for NAB and 31% for βSM-M. Structure prediction from an extended βSM-M sequence based on that alignment led to a high confidence structure prediction closely resembling SARS-CoV-2 βSM-M (RMSD 1.5 Å). However, no other gammacoronavirus showed similar results. Nevertheless, Canada goose coronavirus could serve in further experiments as proof that βSM-M is not completely betacoronavirus-specific as are NAB and the betacoronavirus-specific linker domain.

### Insights of the predicted hexameric assembly of Y1+CoV-Y

The C-terminal region of nsp3, containing the transmembrane domains and the lumenal ectodomain as well as the Y1 and CoV-Y domains, is only partially covered by two PDB entries: 7RQG showing the C-terminal subdomain of CoV-Y (CoV-Yb) [15]; and a larger construct (8F2E) covering Y1b and CoV-Y [16] (Unfortunately, the later study [16] refers to a portion of CoV-Y as “Y3”, which differs from the subdomain labelled “Y3” in the PDB entry 7RQG. To resolve this conflict, we designated these subdomains in Table 2 as “Y1a”, “Y1b”, “CoV-Ya”, and “CoV-Yb”).

Y1 is referred to as “nidovirus-conserved domain of unknown function”, while CoV-Y is a C-terminal nsp3 domain specific to *Coronaviridae* [14]. The lumenal ectodomain was shown to interact with nsp4, resulting in the formation of the viral replication organelles known as “double membrane vesicles” [6]. The cytosolic domains of nsp3 assemble into a hexameric complex participating in the export of new viral RNA-genomes[7].

We analysed the structure prediction of Y1 and CoV-Y and investigated the conservation of Y1 among related viruses. Furthermore, we analysed the prediction of a Y1 hexamer.

#### Structure prediction of Y1 and CoV-Y

For both sarbecoviruses and MHV, a folded structure was predicted by Alphafold2 for the region covering Y1 and CoV-Y (Fig. 7a), consisting of 362 to 377 residues. The predicted model consists of an N-terminal helical loop with low pLDDT, a high confidence fold of ∼160 residues, a short low pLDDT loop, and another high confidence fold (Fig. 7a). The first ∼180 residues correspond to the domain Y1, while the remaining ones are part of CoV-Y.

**Figure 7.**
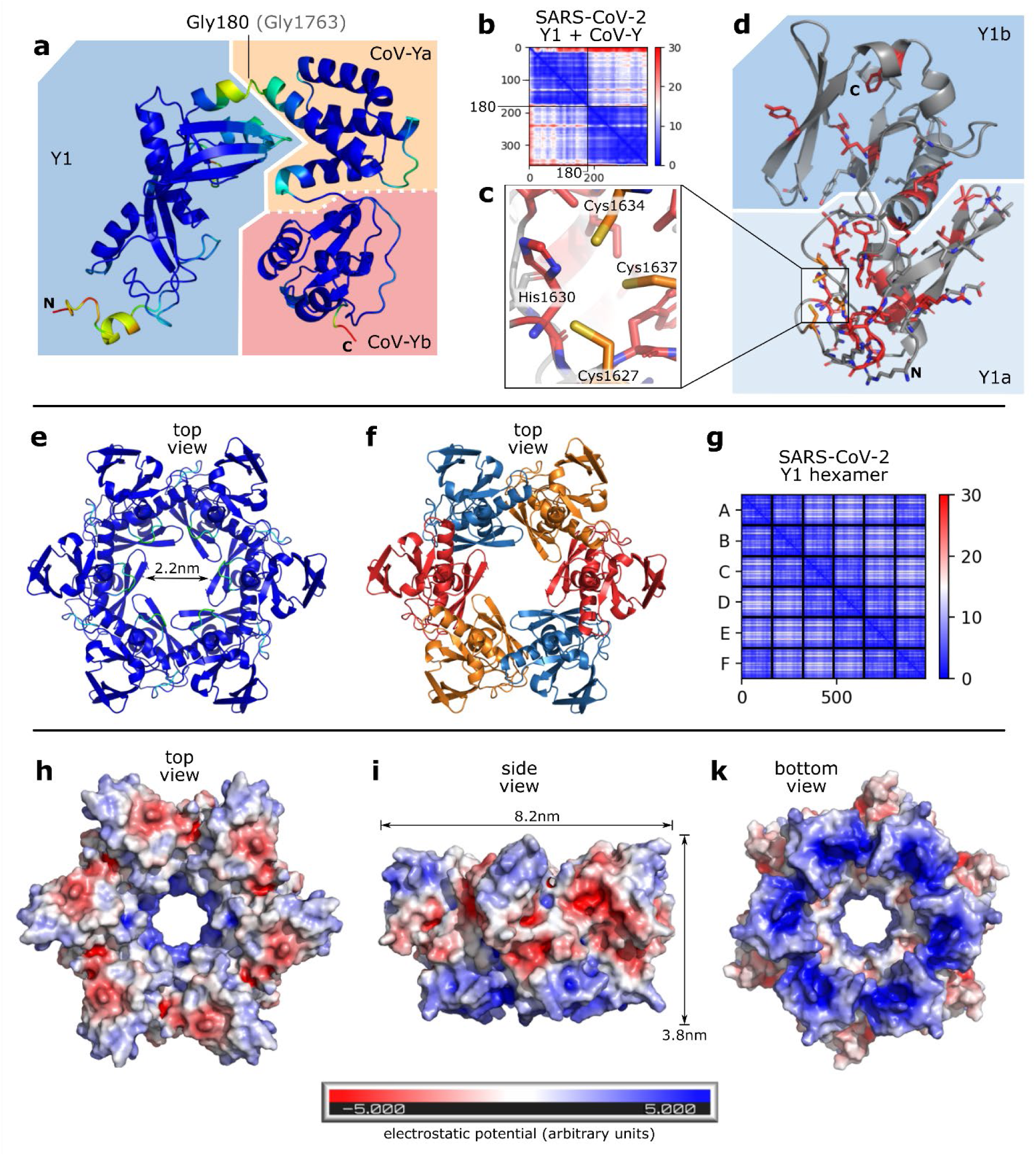
a-b: Prediction of linker+Y1+CoV-Y from SARS-CoV-2 (ranges from Table 2) colored according to pLDDT with blue as high confident (a) with the respective pAE matrix (b). The matrix shows two confident local folds, one with residues before Gly180 and one with residues after it; c: prediction of conserved cysteine-histidine cluster; d: Y1 according to the ranges in Table 2, monomer from hexamer prediction, with amino acids highlighted in red which are conserved among all betacoronaviruses, and the cysteine cluster from (c) in orange. Other shown residues are involved in hexamer formation but are not conserved; e-g: high confidence prediction of Y1 hexamer colored according to pLDDT (e) and by chain (f). The narrowest diameter of the inner gap is 2.2 nm. The pAE matrix shows a highly confident arrangement of all monomers (g); h, i, k: Y1 hexamer with electrostatic surface calculation via PyMOL [30] viewed from different angles. The bottom surface (k) and the surface along the channel (h, k) are positively charged (blue), while the top surface (h) and side surface (i) are mostly neutral (white) or negatively charged (red).

The predicted alignment error (pAE) matrix suggests a clear separation between Y1 and CoV-Y at Gly1763 (SARS-CoV-2 nsp3), but no further division into subdomains (Fig. 7b). However, both domains consist of two distinct globular folds, allowing a separation of the subdomains Y1a and Y1b (Fig. 7d), as well as CoV-Ya, and CoV-Yb (Fig. 7a), which is supported by the experimental structure 8F2E. Only Y1a has not yet been experimentally determined. Repeated predictions of this domain pair show similar results with the same orientation of both domains relative to each other in all cases, but the pAE matrix indicates a potential flexibility in the arrangement and orientation of CoV-Y to Y1.

The fold of Y1a comprises two large β-sheets (21 residues in length), a 50-residues region of loops, and an 11-residue long α-helix (Fig. 7d). This globular fold consisting mostly of loops is held together by numerous hydrogen bonds and contains a conserved cysteine-histidine cluster, which could stabilize the fold by binding metal ions (Fig. 7c).

Y1b makes up the upper half of Y1 (90 residues), containing a triplet of parallel β-sheets, which creates an intertwined structure with a triplet of anti-parallel β-sheets, an α-helix, a loop stabilized by H-bonds, and sharp turns.

The subdomain CoV-Ya consists of four α-helices and CoV-Yb of four α-helices, four β-sheets, and various loops and turns, resulting in a globular fold (Fig. 7a). CoV-Yb resembles closely the PDB structure 7RQG [15] with an RMSD of 0.4 Å. Since Y1a is missing in the PDB entry 8F2E [16], the domains in the prediction are differently packed, resulting in a high RMSD of 2.3 Å. Aligning only the individual subdomains from 8F2E to the prediction in Figure 7a, however, results in RMSD values of 0.4 Å for all three subdomains.

A second potential zinc site, consisting of a histidine and three cysteines was identified in the linker between AH1 (amphipathic helix 1; the domain preceding Y1) [33]. Since the pAE suggests no alignment between N-terminus and the main fold of Y1, it was excluded from the domain boundaries of Y1 and was left as a linker (Table 2). Due to the potential metal-binding cluster, however, it could belong to the amphipathic helix or Y1, which would depend on its function. In SARS-CoV-2 this situation is more complicated, as the linker is shorter and the histidine of the zinc binding site is located in AH1. Without additional information regarding this zinc binding site’s function, it is not possible to define better domain boundaries.

#### Conservation of Y1 among nidoviruses

Despite long sequences of more than 360 residues, the C-terminal domains Y1 and CoV-Y exhibit high sequence similarities between the two sarbecoviruses and MHV, ranging from 65% to 97.5% for Y1 and from 56.5% to 96.5% for CoV-Y (Table 3). Structure predictions show nearly identical folds for Y1 with RMSDs from 0.1 Å to 0.7 Å; CoV-Y shows less similarity with RMSDs from 0.3 Å to 2.4 Å.

Since Y1 is also known as “nidovirus-conserved domain of unknown function” [14], the sequence and structure similarities to other viruses were investigated. First, with other betacoronaviruses, followed by other viruses from *Orthocoronavirinae*, close relatives from *Coronaviridae*, and related nidoviruses.

The global sequence alignment of SARS-CoV-2 with 16 betacoronaviruses (listed in Figure 5), revealed 29 out of 161 amino acids to be conserved across all 17 viruses (Fig. 7d; details in supplementary information); with 22 residues being in Y1a. Further 45 residues of Y1 show conserved chemical properties. The 22 conserved residues in Y1a include a potential zinc binding site [33] made of three cysteines and a histidine (Fig. 7c). With distances of 3.4 Å to 4 Å, the cysteines are too far apart for disulfide bond formation. Furthermore, the conserved amino acids often form hydrogen bonds with each other, which potentially stabilizes the loop-region of Y1a.

Across all examined coronaviruses outside *Betacoronavirus*, Y1 shows moderate sequence similarity from 25.9% to 39.1%, but exhibits high structural similarity with RMSD from 0.6 Å to 1.3 Å in comparison of structure predictions. The closest relatives outside *Orthocoronavirinae* are found within *Pitovirinae*, *Letovirinae*, *Torovirinae, Roniviridae*, *Mesoniviridae*, *Arteriviridae* and *Piscanivirinae* [14]. However, despite showing partially confident fold predictions at sequence identities above 20%, none of the predicted structures resembled the coronaviral Y1, thus questioning Y1’s conservation among nidoviruses (details on used species are in supplementary information).

#### Multimer prediction of Y1 and Y1+CoV-Y

The multimer feature of AlphaFold2 [34] and ColabFold [35] predicts the assembly of homo- and heteromers. It generates a special pAE matrix that evaluates the distances between residues within a monomer and residues of different monomers, thereby assessing the confidence of the predicted complex (Fig. 7g).

Nsp3 is forming a hexameric complex with nsp4 and nsp6 [7]. From all domains, only the prediction of hexameric Y1 (Fig. 7e-f) comes with a confident arrangement of all monomers (Fig. 7g). The average pLDDT of the multimer remains highly confident with a reduction of only 1.4 compared to the monomer. The average pAE per cell of the pAE matrix (Fig. 7g) ranges from 3.2 to 8.4, which is in the confident range of below 10. The low values in the pAE matrix and high pLDDT are consistent across multiple predictions and for all examined viruses. Furthermore, the prediction of the Y1 hexamer and the Y1+CoV-Y hexamer show no Ramachandran outliers.

The hexameric assembly forms a channel with a minimum inner diameter of 2.2 nm. The monomers are held together by several hydrogen bonds, which are formed in SARS-CoV-2 nsp3 by the eleven residues Arg1602, Thr1607, Gly1611, Ser1615, Ala1642, Asp1654, Leu1718, Ser1729, Ser1734, Lys1737, and Leu1746.

Vacuum electrostatics were calculated for the Y1-hexamer (Fig. 7h-k) in PyMOL [30]. These show a primarily positive charge at the inner surface of the channel and on the bottom surface towards the membrane. The sides and the top surface show slightly positive or negative charges areas, as well as some neutral areas and few highly positive charges in buried region.

The multimer prediction of SARS-CoV-2 Y1+CoV-Y leads to a similar complex as solely with Y1. While the Y1 domains remain assembled with high confidence in the pAE matrix, CoV-Y domains show low confidence in the pAE for the alignment between each other and the arrangement to their linked Y1. For MHV, however, the pAE matrix shows much better arrangement, although not such a perfect one as for Y1 in Figure 7g (see Figure S3 of supplementary information).

In UCSF Chimera [36], the Y1-CoV-Y hexamer with lowest pAE values, which was predicted from MHV, was fitted into the cryo electron tomography map from Wolff et al. [7]. This 30.5 Å resolution map shows the whole hexameric pore complex consisting of nsp3, nsp4, and nsp6, and covers also the membrane of the replication organelle. The fitting algorithm finds a position for the model at the base of the pore-complex (Fig. 8). The channel diameter of the model matches that of the map (Fig. 8a-c). From the base, six pillows grow up, were the CoV-Y domains are fitted in. Due to the low resolution, parts of the model are not covered in the volume. Different contour levels change this, but obscure also the channel.

**Figure 8.**
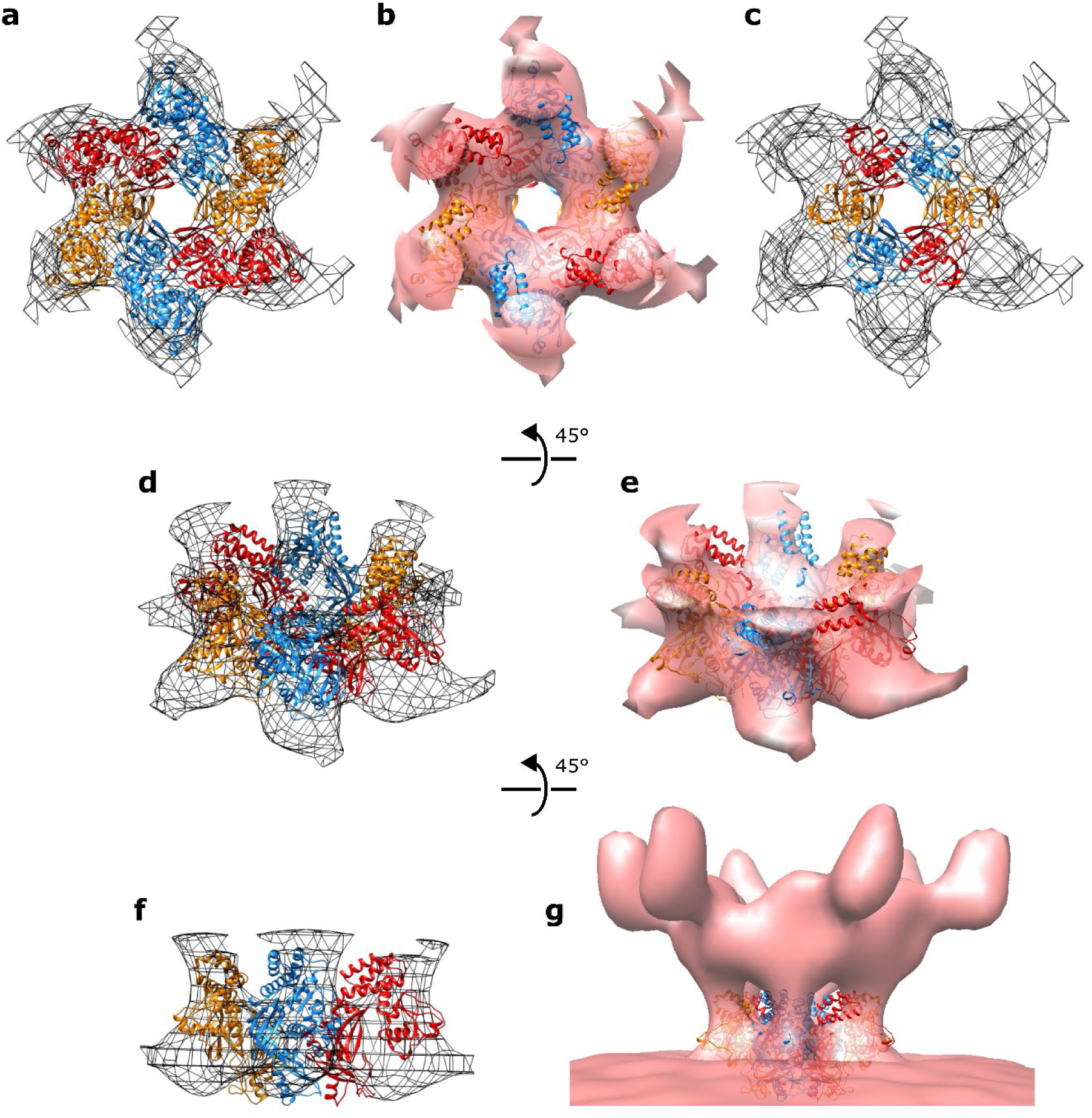
a-b:Top-view of predicted model of Y1+CoV-Y hexamer from MHV fitted into cryo electron tomography map from Wolff et al [7], with volume shown as grid (a) and surface (b). c: same fit, but only Y1 hexamer is shown. d-e: same model-fit as in a-b with rotated camera angle by 45°. f: same model-fit as in d with rotated camera angle by 45°. For more clarity, only half of the hexamer is shown. g: side view of the model fitted into the map with the whole pore-complex visible. All images were generated in UCSF Chimera [36] with a contour level of 2.89. For images a-f, volume was truncated to highlight the complex’s base.

The hexameric prediction of Y1+CoV-Y is not as confident in pLDDT and pAE as the prediction of hexameric Y1, but in the case of MHV, the prediction confidence improves in contrast to the prediction of the Y1+CoV-Y monomer. Despite the uncertainty in the arrangement of both domains (in the pAE matrix), the predicted model fits well into the cryo electron tomography map.

In order to form replication organelles, the ectodomain of nsp3 must interact with nsp4, which is also part of the hexameric complex. Our results revealed a globular fold within the ectodomain and a large ectodomain in nsp4. However, all multimer predictions of the latter domain contradict the experimental evidence and come with low confidence scores.

Expression of a construct containing the Y1 domain was attempted in BL21Gold *E.coli*. However, insufficient amounts of the desired protein were synthesized, preventing an experimental validation of the hexameric Y1 complex. Other, more sophisticated expression hosts (e.g. insect or mammalian cells) might be required to produce useful quantities of Y1.

## Discussion and outlook

Sequence-based structure prediction is transforming the field of structural biology. Although AlphaFold2 predictions are not as reliable as experimental measurements, they can give a good indication of intrinsic disorder and are well suited to assist construct design [26]. We utilized AlphaFold2 to determine domain boundaries more exactly, identified a new domain directly adjacent to the most important nsp3 drug target PL2^pro^, and established a potential mechanism driving the hexameric assembly of the pore.

### Utilizing AlphaFold2 for domain boundary determination and construct design

#### Classification into regions of order and disorder

Correctly predicting secondary structure elements is crucial to differentiate between regions of order and disorder and for the design of crystallizable constructs. Superimposing the AlphaFold2 predictions with experimentally determined domain structures showed that all secondary structure elements were correctly predicted. For all but two, root mean square deviation (RMSD) values were below 1 Å. The two exceptions were Ubl1 from SARS-CoV-1 and MHV with the NMR structures 2GRI and 2M0A, which have a flexible N-terminus. Despite high RMSD values in these cases, the folded region was predicted correctly.

AlphaFold2 predicts disordered loops with low pLDDT values and stretches them away from folded regions, forming the barbed-wire-like conformation first described by Williams et al. [25]. If these disordered regions are large, they can negatively impact the pLDDT of nearby ordered regions and lead to folds differing from the experimentally determined structure, as in the example of the βSM domain. Furthermore, disordered regions with low pLDDT values were sometimes falsely predicted as α-helices. In our cases, such bias could be detected by truncating the low pLDDT loops from the input sequence. This increased the pLDDT of the ordered fold drastically, while the pLDDT of false helices decreased. We also aligned multiple models predicted from the same sequence and observed here good 3D alignment at the ordered sections, while disordered loops and false helices pointed away from the folds in random directions, as illustrated by Figure 3d-e. Transmembrane helices were in our cases also predicted to point away from the main fold in random directions, but they differed from the false helices by having pLDDT values above 90. Since false β-sheets were not observed, the presence of predicted β-sheets could be a good indicator of order.

It is important to note that pLDDT calculation depends on the coverage of the sequence during the multiple sequence alignment step of AlphaFold2 [19]. Sequences with a low number of similar sequences in the database are thus predicted with low pLDDT values. The minimum numbers of similar sequences for ensuring our method to work reliably is yet unclear. In our minimum case, we achieved high pLDDT values (above 80 for majority of residues) with a coverage by 14 sequences with sequence identities from 50% to 100% (for DPUP-like domain from MHV). In a case where sequence identities ranged from 20% to 100% (for SARS-CoV-2 Mac2), sequence coverage of 30 was sufficient.

Conclusively, AlphaFold2 predictions can be used to differentiate between order and disorder by utilizing the pLDDT, presence of predicted secondary structure elements, and the “barbed-wire” phenomenon [25]. It is useful for determining domain ranges of multi-domain proteins and designing crystallizable protein constructs by detecting disordered termini. Shortening the input sequences for AlphaFold2 across multiple runs can improve the confidence measurements and result in structures, which are closer to the experimentally determined structures. Nevertheless, these models remain predictions and are hence only a cheap step in construct design, followed by experimental methods and construction of models based on physical data.

#### Domain boundary determination

In a multidomain protein, experimental structure determination often requires understanding domain boundaries, in order to establish function, internal movement and to solve domains independently. Our method of domain boundary determination is based on AlphaFold2’s capability to predict whether a region is ordered or disordered. During the preparation of short input sequences, covering only few or one domain, we utilized experimentally validated domain boundaries and aligned sequences from related viruses against sequences of established domains. Sequence alignments between homologues already can hint at order or disorder within a multi domain protein, as disordered regions show usually low sequence similarity compared to the similarity of folded domains [37]. This is due to lower selection pressure in disordered regions, which hence accumulate mutations faster.

In our case, we aligned the individual domain sequences between both sarbecoviruses and mouse hepatitis virus (Table S2 of the supplementary information). The folded domains have relatively high sequence similarities, while disordered regions such as linkers had lower similarity. The linker between PL2^pro^ and NAB stood out with a high sequence similarity and was later found to be ordered. We suspect that this betacoronavirus-specific linker domain (further discussed in Section B) went unnoticed due to the incremental annotation of domains for nsp3 in the past. Probably, PL2^pro^ and NAB domains were annotated first and the linker, as all other short linkers in nsp3, was not annotated as such explicitly. Hence, one intuitively assumes this region to be disordered, while in truth it is just not further explored. This case emphasizes the importance of clearly defined ranges and of annotations for disordered regions indicating if they were experimentally proven not to be ordered, such as “unexplored”, “predicted”, or “experimentally validated”.

Our goal was to map each residue in nsp3 to exactly one segment, which can be an individual ordered domain or a region of disorder between domains. Preliminary domain ranges were used as initial input to AlphaFold2. With the increased limit of ColabFold [35] for sequences of up to 4000 residues, predicting the complete structure of a multidomain protein is now also a good starting point for defining domain boundaries without any prior knowledge. In both cases, large regions of disorder can have a negative impact on the prediction of nearby folds as discussed in the previous section. Therefore, the input sequences must be refined by cropping disordered termini. After few iterations, we obtained the final domain ranges for nsp3 (Table 2). This is to our knowledge the first complete list with ranges for each domain and every linker in between of nsp3. However, experiments should be done to validate and complete this list. Until then, it serves as a starting point for construct design and discussion.

### Experimental validation of the Betacoronavirus-specific linker domain (βSLD)

#### Structure analysis

Sequence alignments between SARS-CoV-1 and SARS-CoV-2 showed an unusually high sequence identity for the 34-residue linker located between PL2^pro^, the most important drug target of nsp3, and NAB. Alphafold2 predicted the domain as folded (Figure 4), and further sequence analysis showed this domain to be betacoronavirus-specific, hence we called it betacoronavirus-specific linker domain (βSLD). Among 17 betacoronaviruses we aligned, five out of the 34 residues are conserved (Figure 5) and nine have similar properties [38], suggesting a functional role of this domain. From these 14 residues, 12 are located in a loop or at the edge of a β-sheet right next to a loop (Figure 5). This high selection pressure suggests an important role of the loop residues in function or conservation of the fold.

The betacoronavirus-specific linker domain consists of two β-sheets of two strands each (β1+ β2 and β3+ β4) (Figure 4). Both pairs are interconnected via two hydrogen bonds to each other, forming a compact and plausible fold. The 8-residue central loop is a potential weak point for structural stability but is predicted to form two hydrogen bonds within itself in its backbone. Its flexibility is further limited by a conserved proline, forming a helix-like structure element in the predicted model.

In conclusion, the predicted fold seems valid and structurally stable, although it has one disallowed Ramachandran outlier. Conserved residues overlap with the loop regions where specific residues could be important for maintaining the fold. Lastly, small-angle X-ray scattering (SAXS) measurements agree well with a relaxed model of the predicted structure. Therefore, this domain is very likely forming a stable fold.

#### Potential functions of the linker domain, NAB, and βSM

While PL2^pro^ is conserved across all coronaviruses and beyond, the three following domains, βSLD, NAB, and the betacoronavirus-specific marker domain (βSM), are specific to *Betacoronavirus* [4,14]. The βSM domain comprises an 80-residue long disordered linker (βSM-N) between NAB and the folded part of βSM (βSM-M) as well as another disordered linker βSM-C connecting βSM-M with the transmembrane region.

Since NAB is able to bind DNA and RNA [4], it could support the export of new copies of the viral RNA genome, which would require a position near the channel of the hexameric RNA exporter pore [7]. NAB is connected to the first of two transmembrane domains via βSM and both of its linkers (βSM-N and βSM-C). As Y1 follows the second transmembrane domain and does not have a large linker, its folded part must sit directly adjacent to the pore. Furthermore, the electron tomographic structure expands far into the cytosol [7], requiring most domains to sit on top of each other. Because Mac2, Mac3, DPUP, Ubl2, PL2^pro^, βSLD, and NAB are connected tightly (784 residues in SARS-CoV-2), with only Ubl1 and Mac1 as remaining cytosolic folded domains linked with higher flexibility, all those domains cannot be located adjacent to the pore. This leaves Y1 as basis on the membrane with all remaining domains on top of it as the only plausible option. The βSM domain allows this due to its large disordered regions. In the next section we go into more detail of Y1’s role in this assembly.

To allow interaction between NAB and the viral RNA passing the channel, NAB must be located on top of Y1. However, NAB alone is too small to form a hexameric ring with the same diameter as reported from the cryo electron tomography map [7], so it must assemble into a ring together with additional domains. βSLD and PL2^pro^, as well as potentially βSM-M could fill these gaps and position NAB towards the pore in a functional orientation. In such a scenario, βSLD and βSM-M would serve only a structural purpose. Mutation experiments with disrupted folds of these betacoronavirus-specific domains could give clues about their function in complex assembly and their impact on viral fitness. It is also possible, that βSM-M and βSLD are regulating the activity of NAB, PL2^pro^, and/or other enzymatically active domains which appear close during complex assembly.

#### Evolution of betacoronaviruses-specific domains

Since βSLD, NAB, and βSM are all specific to *Betacoronavirus* [4,14] and potentially interact with each other during complex assembly, their coevolution could hint at their functions. To understand this coevolution, we analysed the gammacoronavirus-specific marker domain (γSM). Only the prediction of Canada Goose Coronavirus γSM resembles the structure of βSM-M, while for all other gammacoronaviruses other folds are predicted. Therefore, the fold of βSM-M is potentially common between both clades in coronaviruses, in contrast to the truly betacoronavirus-specific domains NAB and βSLD. In such a case, the virus would benefit from βSM-M without the presence of NAB or βSLD. However, this statement requires structural validation of the predicted fold of Canada Goose Coronavirus γSM and does not give new clues about the function of βSLD.

Also, we observed high sequence similarities between βSLD from SARS-CoV-2 and another non-structural-protein, endoRNAse, in various alpha-, gamma- and deltacoronaviruses. Structure predictions from these sequences have a similar fold to βSLD but with an additional helix between the central loop and the third β-sheet. A gene-duplication event might have occurred during betacoronavirus evolution, where part of endoRNAse was duplicated and translocated to nsp3. Deletion of the helix-residues allows in our structure prediction a compact and more stable fold. Nevertheless, this gives no information on whether βSLD evolved before, after, or coevolved with NAB, which would be interesting regarding βSLD’s function.

### Insights of the predicted hexameric assembly of Y1+CoV-Y

The cytosolic C-terminal region of nsp3 right after the transmembrane region comprises the domains Y1 and CoV-Y [4,7] (Figure 1b). Only recently, an experimental structure covering the second subdomain of Y1 and the complete CoV-Y domain was published (PDB entry 8F2E) [16]. Sequence analysis combined with structure prediction enabled us to explore the potential function of the “nidovirus-conserved domain of unknown function” during nsp3-complex assembly.

#### Nomenclature

Structure prediction of the complete C-terminal region (residues 1584 to 1945 in SARS-CoV-2 nsp3) shows a high-confidence fold comprising two large domains. Predicted alignment error (pAE) and low pLDDT values around Gly1763 suggest separation into two domains, named Y1 (residues 1599-1759) and CoV-Y (residues 1766-1945), with domain names according to *Lei et al* [4]. Both consists of two closely arranged globular folds and could therefore be divided into subdomains, which must be labelled according to structure and function. Unfortunately, older literature did not define the border between Y1 and CoV-Y, which led to conflict in nomenclature: The PDB structure 7RQG covers the last of four subdomains of Y1+CoV-Y (residues 1844-1945) and is published under the name Y3 (paper not published yet). The second experimentally solved structure from a recent publication (PDB 8F2E) [16] labels this subdomain Y4 and another domain (residues 1764-1847) Y3, making the name Y3 ambiguous.

In the publication of 8F2E [16], the C-terminal region is divided into four domains: Y1 (residue range not given), Y2 (residues 1665-1763), Y3 (residues 1764-1847), and Y4 (residues 1848-1945). These domain ranges closely resemble those from our results, namely Y1a (residues 1599-1664), Y1b (residues 1665-1759), CoV-Ya (residues 1766-1847), and CoV-Yb (residues 1848-1945) respectively. However, the latter study suggests the three latter subdomains to be part of CoV-Y, while we suggest the first two subdomains to be part of Y1 and only the last two as part of CoV-Y, as explained below.

#### Structure prediction of Y1+CoV-Y and domain separation

Domain ranges should generally be based on functional and tightly packed units. In the case of the C-terminal region of nsp3, we therefore consider Y1 with its subdomains Y1a and Y1b as one domain. Both subdomains form a unit in multimer prediction, which gives both subdomains a common function in the composition of the pore complex (see below). Furthermore, conservation of residues of Y1a+Y1b is higher than for the next domain, CoV-Y. Same is true for similarity for predicted structures (Table 3). Cov-Y is also tightly packed, but pAE indicates a clear divide between Y1 and CoV-Y at Gly1763 (see Fig. 7b). While Y1a+Y1b and CoV-Ya+CoV-Yb are packed tightly, Y1 and CoV-Y are connected via a potentially flexible hinge, which is also observable in the PDB structure 8F2E. The fold predictions are also supported by experimental evidence, with an RMSD of 0.4 Å between PDB entry 8F2E and each of the domains Y1b, CoV-Ya, and CoV-Yb, respectively.

For subdomain Y1a, no experimental structure has been solved so far, and our construct comprising Y1a+Y1b was toxic to E. coli during expression (data not included) so that we also were not able to measure it. This is unfortunate, as sequence analysis reveals a high degree of conservation among betacoronaviruses. Most of the conserved residues are in the loop region (residues 1620-1649), which is probably stable due to sidechain interactions. This makes the fold vulnerable to mutations – meaning it must be highly conserved to maintain its fold. Moreover, four of the conserved residues form a cysteine-histidine cluster (three cysteines, one histidine), which is a potential zinc binding site and could stabilize the fold [33].

#### Multimer prediction of Y1 and Y1+CoV-Y

Nsp3 assembles with nsp4 and nsp6 into a hexameric complex, which exports RNA from the replication organelle’s interior into the cytosol [7]. However, the question of what mediates the hexameric assembly of this biologically essential pore remains. We think it may be mediated by the Y1 domain, which becomes clear if considering the spatial arrangement of nsp3. It has two transmembrane domains and both N- and C-terminus, are cytosolic. As Y1 is the first domain on the cytosolic side after the transmembrane region, and since no large linker precedes it, we can assume that this domain forms the foundation of the pore complex right on the membrane. For reasons discussed in the previous section, the domains prior the transmembrane region cannot be located adjacent to the pore, leaving Y1 as the only option.

Our predicted model of Y1+CoV-Y (of which each subdomain agrees well with experimental structures) fits the diameter of the base of the pore (Fig. 8). Because the electron tomographic structure cannot accommodate much more domains at the pore’s base, the N-terminal cytosolic part of nsp3 must go elsewhere, but where exactly? As described in the previous sections, each of the two large linkers preceding the transmembrane region (βSM-N and βSM-C) would be long enough to locate the remaining cytosolic domains on top of the predicted Y1 structure.

Locating Y1+CoV-Y at the base would mean that these domains would need to form a hexameric ring with a channel and in fact, multimer prediction leads to a high confidence hexameric Y1 complex regarding pLDDT and pAE values. Furthermore, the diameter of the pore’s channel of 2.2 nm (Fig. 7e) agrees with the estimates of 2 nm to 3 nm based on the experimental evidence [7]. The monomers are connected through hydrogen bonds between 11 residues in each monomer, of which three residues are conserved among all examined betacoronaviruses and a fourth remains similar in properties [38]. The CoV-Y domains extend away from the channel and are not interacting with each other, leaving Y1 as the only C-terminal domain for holding the assembly together. This is supported by experimental evidence by Li et al., where a construct comprising Y1b+CoV-Y (PDB 8F2E) crystallized in monomeric form [16].

Electrostatics of the predicted Y1 hexamer show a positively charged interior of the pore, which would make it suitable as RNA export channel, as RNA is negatively charged. The bottom surface of the hexamer shows a consistently positive charge, which would allow it to interact with the slightly negatively charged membrane of the replication organelle. Also, the hexamer fits well into the complex’s base of the electron tomographic map (Fig. 8), where membrane contact is unavoidable. Conclusively, we postulate that Y1 forms the foundation of the hexameric pore complex, while the remaining domains are assembling on top of Y1 and CoV-Y, with Y1 being the major contributor in the assembly.

#### Experimental validation

Unfortunately, we were not able to purify Y1 in sufficient quantity to set up crystallization trials. Due to the little published information regarding the purification of the C-terminal domains, and the complete absence of an experimentally determined Y1 structure, we suspect that other researchers were also unsuccessful with this domain. Part of the problem could be the large positively charged surface emerging upon hexamer formation, that could bind to lipid membranes, as well as its potential ability to bind RNA and DNA.

Therefore, we suggest creating mutational variants of Y1 incapable of hexamer formation. Mutating contact residues to alanine might be insufficient, as AlphaFold2 still predicts hexamers with hydrogen bonds to backbone atoms. Mutating to proline, however, works at least in the predictions. Whether effects from mutations are predicted in AlphaFold2 correctly [23,39] or not [40,41] is still to debate. In any case, Y1 mutants with intact fold incapable of hexamer formation can be used for structure solution and for analysis of the hexamer’s impact on the viral fitness, which can finally clarify Y1’s status of a potential drug target.

#### Conservation of Y1 among nidoviruses

Literature labels the Y1 domain as “nidovirus-conserved domain of unknown function”. Furthermore, double membrane vesicles as replication organelles appear in all positive stranded RNA viruses [42], making the need for an RNA-exporter channel present in a large variety of pathogens. Nsp3 has been shown to form an RNA exporter pore in MHV [7], suggesting that nsp3-equivalents in other corona viruses can similarly form exporter-pore-complexes. Conservation in *Nidovirales* supports our previously defined hypothesis of Y1 forming the base of the hexameric RNA exporter channel. In the high-confidence prediction of the Y1 hexamer, multiple hydrogen bonds are predicted, which hold the monomers together. Many of these key residues, as well as an unusually high fraction of all Y1 residues are conserved at least among *Orthocoronavirinae*. However, no similar structure predictions could be generated for viruses outside of *Orthocoronavirinae* and sequence similarities do not rise above 30%, opening the possibility of Y1 being not nidovirus-conserved. In such a case, related proteins with different folds as well as alternative mechanisms for pore-assembly are thinkable.

Conclusively, our findings suggest Y1 to be a coronavirus-conserved domain likely involved in pore-complex assembly. Due to its high conservation in *Orthocoronavirinae*, it is not only an interesting drug target in human-infecting betacoronaviruses, but also in gammacoronaviruses, which infect various bird species and are therefore a potential threat for livestock. Researching this domain could thus lead to therapeutics targeting a wide range of viral diseases.

### Conclusion and outlook

Non-structural protein 3 is a large multi-domain protein from SARS-CoV-2 with two established drug target domains and several domains with unresolved functions or structure. Its long evolutionary history and presence across distantly related viruses point at its vital function to the infection cycle, of which we may have uncovered only a fraction. Our discovery of the betacoronavirus-specific linker domain underlines this aspect and demonstrates the need for better domain annotation. Information about the exploration status, such as “unexplored”, “predicted”, or “experimentally validated” would improve our understanding of multi-domain proteins and highlight knowledge gaps. We demonstrated that automated use of modern structure prediction is an excellent starting point for domain annotation in large multi-domain proteins and can also provide constructs for experimental validation.

Electron tomography of NSP3 shows that it forms a large hexameric pore for the export of RNA into the cytosol. Based on high confidence predictions we put forward the hypothesis that the Y1 domain drives the hexameric assembly. This is supported by conservation of key residues regarding assembly and folding, electrostatics, fitting the multimeric model into the experimental electron tomography map, and by geometrical consideration regarding domain vicinity in the assembled complex. If purification of Y1 is successful, validation of the hexameric assembly by size exclusion is a logical next step. Evaluation of Y1’s impact on the total number of pore-complexes via mutation experiments will determine Y1’s influence on viral fitness and clear its status as a potential drug target.

Research into the interaction between Y1 and other domains, such as NAB, βSM-M, and the betacoronavirus-specific linker domain, which could bind to the top part of Y1+CoV-Y, as well as the binding of Y1-hexamer to lipid membranes may shed more light on NSP3 and the biological mechanisms of coronaviruses.

## Methods

### Utilizing AlphaFold2 for domain boundary determination and construct design

We used the free access version of AlphaFold2 via Google Colab, ColabFold [35], with default settings (without templates and without relaxation), using the MMSeqs2 algorithm [43] for multiple sequence alignment. For both sarbecoviruses, we used the version v2.1.1, while for MHV, we used v2.1.2. Towards the end of our work, v2.3.1 was released, which was used for submission of full length nsp3 sequences due to a larger input sequence limit.

Protein sequences of SARS-CoV-2, SARS-CoV-1, and MHV nsp3 were obtained from NCBI (reference ids YP_009742610.1, NP_828862.2, and NC_048217.1, respectively).

The preliminary domain ranges for all nsp3 domains of SARS-CoV-1 were defined first, since it had the most experimentally determined domain structures in the PDB [44]. The solved domains included Ubl1, Mac1, Mac2, Mac3, DPUP, Ubl2, PL2^pro^, and NAB. For prediction of transmembrane domains the full nsp3 sequence was submitted to the TMHMM 2.0 server [27]. Domain ranges of disordered domains or unresolved domains, i.e. proposed domains not associated with experimentally determined structures, were taken from literature [4]. The remaining regions between all domains were designated as linker regions.

The preliminary domain ranges of SARS-CoV-2 and MHV were defined by an analogous procedure with the respective domain structures from the PDB, the gene annotations in the NCBI entry, transmembrane domain prediction, and literature research. For SARS-CoV-2, the PDB [44] contained at the time of this work structures for the domains Ubl1, Mac1, Ubl2, PL2^pro^, NAB, part of βSM, and CoV-Yb. At the end of our work, the domains Y1b and complete CoV-Y were published, but not used for our initial domain determination. For MHV, PDB contained Ubl1, the DPUP-like domain, Ubl2, and PL2^pro^. To define the ranges of the remaining domains in SARS-CoV-2 and MHV, we performed global sequence alignments with Clustal Omega [45] between the preliminary ranges of SARS-CoV-1 and the full nsp3 sequence of the other two viruses. Each residue of all three nsp3 sequences was assigned to a domain or a linker between domains, respectively.

The sequences of each preliminary domain range were submitted to ColabFold [35]. In order to evaluate the prediction’s accuracy, we aligned the models predicted by AlphaFold2 with the respective experimentally determined structures from the PDB [44] in PyMOL [30] and calculated there the root mean square deviation (Table 1).

The AlphaFold2 results and their pLDDT values were utilized to determine regions of order and disorder for all submitted sequences. The classification process is outlined in Figure 2. Regions with a lack of secondary structure elements and average pLDDT values below 50 were considered disordered, where secondary structure elements such as α-helices and β-sheets were recognized manually from cartoon representation in PyMOL [30], which were recognized by PyMOL’s internal DSSP algorithm. These included also so-called “barbed-wire” regions with no proper torsion angles or hydrogen bonds as described by Williams et al. [25]. Secondary structure elements with pLDDT above 80 and loops with pLDDT 50-80 flanked by secondary structure elements were considered ordered.

Residues in secondary structure elements (α-helices and β-sheets) with pLDDT values below 80 were evaluated individually. First, we truncated the potentially disordered regions at the termini and resubmitted the shortened sequence, which resulted in different pLDDT values. If the pLDDT of a region increased drastically (values rising from below 50 to over 80), the region was considered ordered henceforth. Regions containing α-helices where pLDDT stayed below 80 after cropping, and which pointed away from the main fold, were classified as potentially disordered, especially if the pLDDT lowered after cleavage. Regions containing β-sheets were classified as ordered.

Structure predictions containing large proportions of low pLDDT (such as βSM domain) were iteratively truncated and resubmitted, resulting in increasing pLDDT values for the folded regions. Iterative truncation of all domains was performed at the termini, if more than three terminal residues showed pLDDT values below 50. At each iteration, we cleaved regions we classified as disordered (via the method above) completely except for three residues at the border where the pLDDT rises above 50.

In addition to the individual domain ranges, we submitted sequences to AlphaFold2 which covered multiple domains and linkers (see Figure S1 of the supplementary information). These predictions contained two or more consecutive domains positioned relative to each other and the predicted Alignment Error (pAE) was used to assess the validity of such assemblies. Consistent values close to zero in the pAE matrix across all residues were considered as plausible alignment between two domains. Average pAE values above 15 were considered as implausible. Such alignment information was utilized to identify subdomains, which are part of a larger domain. To calculate average pLDDT and pAE values, we used a custom python script. All results together were considered for defining the final ranges of each individual domain, where the pAE was used last.

Ramachandran outliers for the predicted folds of the betacoronavirus-specific linker domain and the Y1 hexamer were calculated by MolProbity [31] in order to evaluate the prediction’s validity.

### Experimental validation of the Betacoronavirus-specific linker domain (βSLD)

Using the method described before, we identified a new folded domain between PL2^pro^ and NAB (residues 1057-1090 in SARS-CoV-2 nsp3), which was analysed further with bioinformatical and experimental methods.

#### Analysis of predicted structure and conservation of its sequence

The new fold was checked visually in PyMOL [30], with a focus on hydrogen bonds by applying the preset “technical”. The sequence of the new domain (residues 1057-1090 in SARS-CoV-2 nsp3) was submitted to BLAST [46], with SARS-CoV-2 sequences excluded. In additional BLAST queries, the whole taxonomy of *Coronaviridae* was excluded to find other homologs. Furthermore, we used the sequences of all betacoronaviruses found under the NCBI taxonomy id 694002 in local pairwise sequence alignments against the sequence of the new domain from SARS-CoV-2. The alignment was performed via EMBOSS Water version 6.6.0 [45]. Target sequences from other betacoronaviruses were annotated equivalents of SARS-CoV-2 nsp3, if available. Otherwise, the sequence of the respective polyprotein 1ab was used, which contains all nsps. In the latter case, we ensured that the alignment was potentially localized within the nsp3 region on the polyprotein. The sequences from the alignment results were extended to match the length of the domain in SARS-CoV-2 and were afterwards submitted to ColabFold. The whole procedure was repeated for the remaining *Coronaviridae*.

Since the two domains following the linker domain, NAB and βSM domain, were previously assumed to be betacoronavirus-specific based on sequence information alone [14], we applied the same approach on those two domains as well.

#### Expression of NSP3 PL2^pro^+LinkerDomain

To validate the presence of the discovered linker domain, *in vitro* experiments were performed. The construct containing the domain Ubl2, PL2^pro^ and the linker domain (SARS-CoV-2, Wuhan original strain; C111S mutant; see supplementary information section B) was amplified by PCR from a template kindly provided by David LV Bauer (Francis Crick Institute). The coding sequence was ligated into a pGEX-6P vector using (5’) BamHI & NotI (3’) restriction sites introduced during amplification. Correct insertion was confirmed by Sanger sequencing. Recombinant protein was expressed in LB in BL21Gold *E.coli* as a GST-3C-fusion protein. After induction at OD_600_ of 0.6, the temperature was reduced to 16°C, and cells were harvested the following morning. Cell pellets were frozen at -80°C until needed.

#### Purification of NSP3 PL2^pro^+LinkerDomain

Cell pellets were resuspended in lysis buffer (50mM Tris pH 7.5, 300mM NaCl, 5% (v/v) Glycerol, 0.5mM TCEP, 1µM Zinc Acetate. EDTA-free protease inhibitors (Roche) were added as per manufacturer’s instructions. Cells were lysed by sonification, and lysate clarified by centrifugation at 45,000*g*, 4°C for 45 minutes. Protein was harvested by incubating the lysate with Glutathione Sepharose 4B (Cytiva) for 2 hours at 4°C with constant mixing. The beads were then harvested and washed with 10 bed-volumes of lysis buffer and then 20 bed-volumes of SEC buffer (20mM Tris, pH 7.5, 150mM NaCl, 0.5mM TCEP, 1uM Zinc Acetate). Beads were then resuspended in 5 bead volumes of wash buffer, and then incubated with GST-HRV3C protease overnight at 4°C with constant mixing. The cleaved product was collected from the beads via filtration. This material was then concentrated to 5mg/mL, aliquoted, snap frozen in liquid nitrogen and stored at -80°C until needed.

#### Crystallization of NSP3 PL2^pro^+LinkerDomain

Prior to crystallization, the protein was thawed on ice and any remaining aggregates or impurities were removed via a final size exclusion chromatography step, using a Superdex200 increase column, equilibrated in SEC buffer (20mM Tris-HCl pH 7.5, 150mM NaCl, 1mM TCEP, 1µM zinc acetate). This material was diluted 1:2 with milliQ water and concentrated to 7.5mg/mL for crystallization trials. Sitting-drop vapour diffusion crystallisation experiments were setup in MRC 2-well 96 well plates using a Formulatrix NT-8 drop setting robot. Initial microcrystals we obtained using small (200nl protein + 100 mother liquor) drops, which were then used to streak seed into larger 400nl + 200nl drops. Crystals appeared after ∼4-5 weeks after incubation & streak seeding at 4°C in 0.1M MES pH 6.8, 8.4% PEG20K. Crystals were cryocooled in liquid nitrogen using crystallization liquor supplemented with 20% Ethylene glycol.

Diffraction data were collected at beamline I24 at Diamond Light Source. Auto processing using the AutoProc pipeline [47] indicated that the data extended to ∼2.1 Å resolution, and that the crystals had space group P1. Molecular replacement using the MRBUMP pipeline [48] in CCP4Cloud [49] gave a reasonable solution using PDB 4M0W (“Crystal Structure of SARS-CoV papain-like protease C112S mutant in complex with ubiquitin” [50]) with a TFZ of 12.0 and an initial *R*_free_ of 47%.

Inspection of the electron density map in COOT [51] revealed that the amino-terminal lobe of PL2^pro^ was well defined, but the carboxy-terminal lobe was not, with poor electron density and noisy difference maps. The output from this molecular replacement solution was submitted to the Modelcraft auto-building & refinement pipeline [52] which finished with a final *R*_free_ of 26.5%. Inspection of the results revealed that the crystals in fact contained two copies of the N-terminal lobe (Arg3 to Cys181 (construct numbering, supplementary information section B)) of the PL2^pro^ monomer, forming a close and compact dimer. No further refinement was undertaken.

#### SAXS analysis of NSP3 PL2^pro^+LinkerDomain

The purified protein was shipped on dry ice to P12 BioSAXS beamline [53] at PETRA III (DESY, Hamburg, Germany), where the small-angle X-ray scattering (SAXS) experiment was conducted. P12 provides monochromatic X-rays with a 0.2 x 0.05 mm^2^ beam at the sample capillary. For this experiment, the X-ray wavelength and sample-to-detector distance were 0.124 nm and 3000 mm respectively. The purified monodisperse fraction was prepared as protein stocks with a concentration of 15 mg/mL. The stock buffer is 20 mM Tris (pH 8.5), 150 mM NaCl, 0.5 mM TCEP, 1μM zinc acetate. The buffer for background subtraction was prepared by first diluting the protein stock 1:1 (v:v) with a close match of the stock buffer then collecting the flow-through of ultrafiltration when the diluted protein solution was concentrated to the original volume. The concentration series (0.65 mg/mL, 0.97 mg/mL, 1.29 mg/mL, 1.61 mg/mL, 1.94 mg/mL, 3.22 mg/mL) is prepared by dilution with the closely matched buffer and careful filtration with Nanosep 100K OMEGA filters (PALL Life Scienes). The concentration series was measured under the standard “batch mode” at P12. The scattering profile was radially integrated, averaged and absolutely scaled from the corresponding detector images of 30 exposures. Each exposure was 95 ms. The model and its theoretical SAXS curve are from the most fitted output generated by SREFLEX [32], using the background subtracted SAXS profile of the 3.22 mg/mL sample and relaxed Alphafold2 prediction as the input. The ab-initio modeling was carried out as following: 1) generate 20 bead models with DAMMIF [54] using the GNOM [55] output of the 3.22 mg/mL sample; 2) generate a starting search volume by averaging the 20 bead models using DAMAVER [56]; 3) run DAMMIN [57] using the starting search volume and the GNOM output of the 3.22 mg/mL sample.

### Insights of the predicted hexameric assembly of Y1+CoV-Y

#### Conservation of Y1 among Nidovirales

After determining new domain ranges for the Y1 domain, the nidovirus-conserved domain of unknown function, a protein-protein BLAST search [46] with Y1 from SARS-CoV-2, excluding SARS-CoV-2, was performed. Afterwards, global sequence alignments with this sequence and the polyprotein sequences of other viruses were performed with Clustal Omega on default settings [45] to identify sequences of Y1-equivalents. These sequences were then submitted to AlphaFold2 [19] via ColabFold [35]. The predicted structures were aligned in PyMOL [30] to the structure prediction of Y1 from SARS-CoV-2 in order to calculate the root mean square deviation (RMSD) to measure the predicted structural similarity. The examined viruses are listed in the supplementary information section B.

#### Multimer prediction of Y1+CoV-Y

Based on the hexameric assembly of nsp3 in complex with nsp4 and nsp6, which was observed by cryo electron tomography [7], and Y1’s potential location at the pore’s base, a hexamer prediction of the Y1 domain was performed via AlphaFold2’s multimer feature [34]. It was afterwards repeated for each other domain. Since the prediction did not complete for large domains at default settings, the code in Colab was adapted to output only one model by setting “num_models” to 1 and model_order to “[1]”. The confidence of multimeric arrangements was evaluated with the pAE matrix, were average pAE values below 10 were handled as confident. Electrostatics of confidently predicted hexamers were calculated in PyMOL [30] via “generate vacuum electrostatics”.

Fitting the models to the cryo electron tomography density map was accomplished in UCSF Chimera [36], by first positioning it manually at the base of the pore-complex in the correct orientation and afterwards applying Chimera’s fitting algorithm under Tools, Volume Data, Fit in Map. Rotation and shift were allowed.

#### Expression and Purification of Y1

The construct of Y1 (supplementary information section C) was amplified by PCR from a template kindly provided by David LV Bauer (Francis Crick Institute). The coding sequence was ligated into a pGEX-6P vector using (5’) BamHI & NotI (3’) restriction sites introduced during amplification. Correct insertion was confirmed by Sanger sequencing. Recombinant protein was expressed in LB in BL21Gold *E.coli* as a GST-3C-fusion protein. After induction at OD_600_ of 0.6, the temperature was reduced to 16°C, and cells were harvested the following morning. Cell pellets were frozen at -80°C until needed.

Cell pellets were resuspended in lysis buffer (50mM Tris pH 7.5, 300mM NaCl, 5% (v/v) Glycerol, 0.5mM TCEP, 1µM Zinc Acetate. EDTA-free protease inhibitors (Roche) were added as per manufacturer’s instructions. Cells were lysed by sonification, and lysate clarified by centrifugation at 45,000*g*, 4°C for 45 minutes. Protein was harvested by incubating the lysate with Glutathione Sepharose 4B (Cytiva) for 2 hours at 4°C with constant mixing. The beads were then harvested and washed with 10 bed-volumes of lysis buffer and then 20 bed-volumes of SEC buffer (20mM Tris, pH 7.5, 150mM NaCl, 0.5mM TCEP, 1uM Zinc Acetate). Beads were then resuspended in 5 bead volumes of wash buffer, and then incubated with GST-HRV3C protease overnight at 4°C with constant mixing. Analysis by SDS-PAGE of the purification process showed that no detectable Y1 had been expressed.

## Acknowledgements

Thanks to Gianluca Santoni, Lea von Soosten, the Coronavirus Structural Taskforce [17], Arwen Pearson, Andrew Torda, and Kay Grünewald for interesting and insightful discussions.

Huge thanks goes to David LV Bauer from Francis Crick Institute, who provided us the template for our PL2^pro^+linker_domain construct.

We thank Diamond Light Source for beamtime (proposal mx25587) and the involved staff at I24 for assisting with data collection.

We acknowledge EMBL Hamburg (Desy, Hamburg, Germany) for kindly providing the emergency beamtime. We would like to give special thanks to Cy Jeffries (EMBL, Hamburg) for his assistance over the course of the BioSAXS experiments. The beamtime was allocated as a part of the SPC-CSSB BAG proposal.

This work was supported by the German Federal Ministry of Education and Research [grant no. 05K19WWA and 05K22GU5] and Deutsche Forschungsgemeinschaft [project TH2135/2-1]. D.C.B. acknowledges that this work was supported by the Francis Crick Institute, which receives its core funding from Cancer Research UK (CC2068), the UK Medical Research Council (CC2068) and the Wellcome Trust (CC2068)

## Supplementary Information

### Utilizing AlphaFold2 for domain boundary determination and construct design

#### Definition of preliminary domain boundaries

During the process of our research, several domain structures became experimentally solved, so the following procedure started with less information available and newer data was utilized later during the process. For determination of preliminary domain ranges, which later served as input sequences for AlphaFold2 [19], we started with SARS-CoV-1, as it had the highest number of solved domains and was augmented by theoretical domain ranges from Lei et al [4]. Our goal was to map every residue of the SARS-CoV-1 nsp3 sequence (NCBI number NP_828862.2) to exactly one domain. We started with available experimental structures, which covered the domains **Ubl1** (PDB codes 2GRI, 2IDY), **Mac1** (2FAV, 2ACF), **Mac2** (6YXJ), **Mac3** (2JZE, 2JZD, 2JZF, 2RNK), **DPUP** (2KAF, 2KQW), **Ubl2+PL2^pro^** (4M0W, 5TL6, 3E9S, 4OVZ, 3MJ5, 5Y3Q, 2FE8, 4OW0, 5Y3E, 5E6J, 4MM3, 5TL7), and **NAB** (2K87). Structures 2W2G and 2WCT include the sequence of Mac2 and Mac3.

From the available Mac1 structures, it was unclear whether residues 356-358 (SARS-CoV-1 nsp3 numbering) belong to the linker following Mac1, or if these residues and parts of the linker must be included in Mac1. Unclear regions were resolved later with data available from AlphaFold2.

For Mac2 and Mac3 it was unclear whether residues 513-526 belong to the C-term of Mac2 or to the N-term of Mac3, since the structures 2JZE, 2JZD, 2JZF, 2RNK show both cases. However, 2KQV (from other authors) suggests Mac3 to begin at Gly527. The structure 6YXJ (from a third group of independent authors) supports this domain separation, as Mac2 ends here with residue Leu526. The last ten residues, however, are not modelled, hence we defined the preliminary range of Mac2 to end at Ser516, while Glu517 to Leu526 were defined as short linker.

By knowing the domain ranges of Ubl1 and Mac1 based on experimental structures, HVR was defined automatically by the range in between. Same rule applies to DPUP domain surrounded by Mac3 and Ubl2+PL2^pro^.

While all mentioned domain ranges were defined by structural information, all remaining ranges were taken from Lei et al. [4]. After mapping each residue of SARS-CoV-1 nsp3 to a domain or linker, global sequence alignments were performed between each of those preliminary domain ranges (Table S1, SARS-CoV-1 column) and full length nsp3 of SARS-CoV-2 or MHV. Preliminary domain ranges for SARS-CoV-2 and MHV were based on these alignment results, but ranges based on experimental structures were prioritized, if available.

For SARS-CoV-2, available structures covered the domains **Ubl1** (PDB code 7KAG), **Mac1** (used PDB codes: 6WEY, 6WOJ, 7CZ4, 6YWL, 7BF5, 7KQP), **Ubl2+PL2^pro^** (7CMD, 7CJD, 7CJM, 7LLZ), **NAB** (7LGO), and **CoV-Yb** (7RQG). For Mac1 and Ubl2+PL2^pro^ a great number of structures was available. The listed ones are the few we used to define the domain ranges. Towards the end of our research, the structure of **βSM** (7T9W) was published, which validated our results of a folded central region surrounded by two disordered linkers. For MHV, experimental structures provided the ranges of **Ubl1** (2M0A), **DPUP-like** (4YPT), and **Ubl2+PL2^pro^** (5WFI). The ranges from Lei et al. were used for the remaining ranges.

Since most of the utilized structures were deposited in the PDB after 2018, our ranges deviate from those by Lei et al. [4]. Both, new and old ranges, are listed in Table S1.

**Table S1.** Preliminary residue ranges of nsp3 domains from SARS-CoV-2, SARS-CoV-1, and MHV in comparison to ranges defined in previous literature. Previously defined ranges for SARS-CoV-1 and MHV are from Lei et al. [4], whereas previous ranges for SARS-CoV-2 nsp3 are from NCBI gene annotations with the reference id YP_009742610.1. a: These ranges were not stated explicitly and are derived from the surrounding ranges. b: These ranges had no defined start/end. c: These ranges were predicted by TMHMM 2.0. d: This domain was not determined in the gene annotations. Ranges with a yellow shade are identical between the preliminary and the previous ranges. Ranges with a blue shade differ by a maximum of 5 residues into each direction.

| Complete Name | Preliminary residue ranges for SARS-CoV-2 | Previously defined residue ranges for SARS-CoV-2 | Preliminary residue ranges for SARS-CoV-1 | Previously defined residue ranges for SARS-CoV-1 | Preliminary residue ranges for MHV | Previously defined residue ranges for MHV |
| --- | --- | --- | --- | --- | --- | --- |
| Ubiquitin-like domain 1 (Ubl1) | 1-111 | ???<sup>d</sup> | 1-112 | 1-112 | 1-115 | 20-109 |
| Hypervariable region (HVR) | 112-206 | ???<sup>d</sup> | 113-181 | 113-183 | 116-252 | 113-243 |
| Papain-like protease 1 (PL1<sup>pro</sup>) | - | - | - | - | 253-480 | 252-500 |
| Macrodomain 1 (Mac1) | 207-379 | 234-359 | 182-358 | 184-365 | 481-655 | 509-633 |
| Linker Mac1-Mac2/Linker Mac1-DPUlike | 380-412 | 360-414<sup>a</sup> | 359-388 | 366-388<sup>a</sup> | 656-693 | 634-701<sup>a</sup> |
| Macrodomain 2 (Mac2) | 413-550 | 415-540 | 389-516 | 389-524 | - | - |
| Linker Mac2-Mac3 | - | - | 517-526 | - | - | - |
| Macrodomain 3 (Mac3) | 551-675 | 533-675 | 527-651 | 525-652 | - | - |
| Domain preceding Ubl2 and PL2<sup>pro</sup> (DPUP) | 676-745 | 678-743 | 652-722 | 653-720 | 694-777 | 702-776 |
| Ubiquitin-like domain 2 (Ubl2) | 746-804 | 748-???<sup>b</sup> | 723-781 | 723-???<sup>b</sup> | 778-837 | 777-???<sup>b</sup> |
| Papain-like protease 2 (PL2<sup>pro</sup>) | 805-1063 | ???-1050<sup>b</sup> | 782-1040 | ???-1036<sup>b</sup> | 838-1080 | ???-1074<sup>b</sup> |
| Linker PL2<sup>pro</sup>-NAB | 1064-1088 | 1051-1994<sup>a</sup> | 1041-1065 | 1037-1065<sup>a</sup> | 1081-1116 | 1075-1110<sup>a</sup> |
| Nucleic-acidic-binding domain (NAB) | 1089-1203 | 1095-1201 | 1066-1180 | 1066-1180 | 1117-1232 | 1111-1229 |
| Betacoronavirus-specific marker domain (βSM) | 1204-1412 | 1226-1341 | 1181-1390 | 1203-1318 | 1233-1448 | 1277-1401 |
| Transmembrane domain 1 (TM1) | 1413-1435<sup>c</sup> | 1414-1435 | 1391-1413<sup>c</sup> | 1391-1413<sup>c</sup> | 1449-1471 | 1450-1471 |
| Nsp3 ectodomain | 1436-1531<sup>c</sup> | 1436-1518<sup>a</sup> | 1414-1495<sup>c</sup> | 1414-1495<sup>c</sup> | 1498-1564 | 1472-1577<sup>a</sup> |
| Transmembrane domain 2 (TM2) | 1532-1554<sup>c</sup> | 1519-1541 | 1496-1518<sup>c</sup> | 1496-1518<sup>c</sup> | 1565-1587 | 1578-1600 |
| Linker TM2-AH1 | 1555-1560<sup>c</sup> | ???<sup>d</sup> | 1519-1522<sup>c</sup> | 1519-1522<sup>a, c</sup> | 1588-1607 | ???<sup>d</sup> |
| Amphipathic helix 1 (AH1) | 1561-1583<sup>c</sup> | ???<sup>d</sup> | 1523-1545<sup>c</sup> | 1523-1545<sup>c</sup> | 1608-1630 | ???<sup>d</sup> |
| Nidovirus-conserved domain of unknown function (Y1) | 1584-???<sup>b</sup> | ???<sup>d</sup> | 1546-???<sup>b</sup> | 1546-???<sup>b</sup> | 1631-???<sup>c</sup> | ???<sup>d</sup> |
| Coronavirus-specific C-terminal domain (Y2) | ???<sup>b</sup>-1843 | ???<sup>d</sup> | ???-1820 | ???-???<sup>b</sup> | ???-1904 | ???<sup>d</sup> |
| Coronavirus-specific C-terminal domain (Y3) | 1844-1945 | ???<sup>d</sup>-1944 | 1821-1922 | ???-1922<sup>b</sup> | 1905-2006 | ???-2004 |

#### Sequence comparison of domains between SARS-CoV-2, SARS-CoV-1, and MHV

Sequences based on the preliminary domain ranges were compared in pairwise sequence alignments (Table S2). It is noteworthy that all regiones expected to be disordered (by being a linker or explicitly stated as disordered in Lei et al. [4]) show relatively low values, except for the linker between PL2^pro^ and NAB. Furthermore, the C-terminal domains Y1 and CoV-Y show despite their large size very high similarity.

**Table S2.** Results from local pairwise sequence alignments with domain sequences from preliminary ranges (stated in Table S1). The alignments were calculated via EMBOSS Water version 6.6.0 [45]. Domains not present in a virus are marked with “-“. “x” indicates a failed alignment (less than 10 residues aligned). Blue indicates domains assumed in literature to have a stable fold; red indicates linkers and domains assumed to be mostly disordered; yellow indicates domains in the transmembrane region.

| Domain name | SARS-CoV-2 / SARS-CoV-1 |  | SARS-CoV-2 / MHV |  | SARS-CoV-1 / MHV |  |
| --- | --- | --- | --- | --- | --- | --- |
|  | Sequence identity | Sequence similarity | Sequence identity | Sequence similarity | Sequence identity | Sequence similarity |
| Ubiquitin-like domain 1 (Ub1) | 78.6% | 89.3% | 38.3% | 52.2% | 35.4% | 53.1% |
| Hypervariable region (HVR) | 43.8% | 64.0% | 21.7% | 38.7% | 25.6% | 35.4% |
| Macrodomain 1 (Mac1) | 73.8% | 85.1% | 32.5% | 52.6% | 34.9% | 51.7% |
| Linker Mac1-Mac2 | 41.4% | 51.7% | 38.5% | 69.2% | x | x |
| Macrodomain 2 (Mac2) | 71.1% | 85.2% | - | - | - | - |
| Macrodomain 3 (Mac3) | 81.6% | 89.6% | - | - | - | - |
| Domain preceding Ubl2 and PL2<sup>pro</sup> (DPUP) | 73% | 87% | 25% | 50% | 40% | 60% |
| Ubiquitin-like domain 2 (Ubl2) | 90% | 97% | 31% | 54% | 33% | 52% |
| Papain-like protease 2 (PL2<sup>pro</sup>) | 81.1% | 89.6% | 32.6% | 47.1% | 35.1% | 48.3% |
| Linker PL2<sup>pro</sup>-NAB | 80% | 88% | 40% | 48% | 36% | 48% |
| Nucleic-acidic-binding domain (NAB) | 81.7% | 89.6% | 32.0% | 44.0% | 29.0% | 42.0% |
| Betacoronavirus-specific marker domain (βSM) | 69.2% | 82.2% | 20.3% | 37.7% | 22.8% | 39.2% |
| Transmembrane domain 1 (TM1) | 77.0% | 82.0% | x | x | x | x |
| Nsp3 ectodomain | 71.0% | 82.0% | 32.0% | 51.0% | 38.0% | 52.0% |
| Transmembrane domain 2 (TM2) | 80% | 80% | x | x | x | x |
| Amphipathic helix 1 (AH1) | x | x | 29.0% | 48.0% | 29.0% | 53.0% |
| Nidovirus-conserved domain of unknown function (Y1) + Coronavirus-specific C-terminal domain a (CoV-Ya) | 88.1% | 95.4% | 37.7% | 62.3% | 37.9% | 61.7% |
| Coronavirus-specific C-terminal domain b (CoV-Yb) | 90.2% | 100.0% | 40.6% | 58.4% | 39.6% | 58.4% |
| full nsp3 sequence | 76.0% | 86.5% | 23.6% | 37.4% | 23.4% | 37.2% |

#### Domain separation via predicted Alignment Error

The sequences from preliminary domain ranges were submitted to AlphaFold2 [19]. The main text explains how the results were processed and how to differentiate between ordered and disordered regions.

To get an impression of the relative alignment of adjacent domains, nine multi-domain sequences in SARS-CoV-2 nsp3 were submitted to Alphafold2. The pAE matrices of the rank1 prediction from each case are depicted in Figure S1. Overlap with the transmembrane region was performed only in the full length nsp3 prediction, since AlphaFold2 does not take membranes into account, leading to biologically incorrect arrangements of domains.

**Figure S1.**
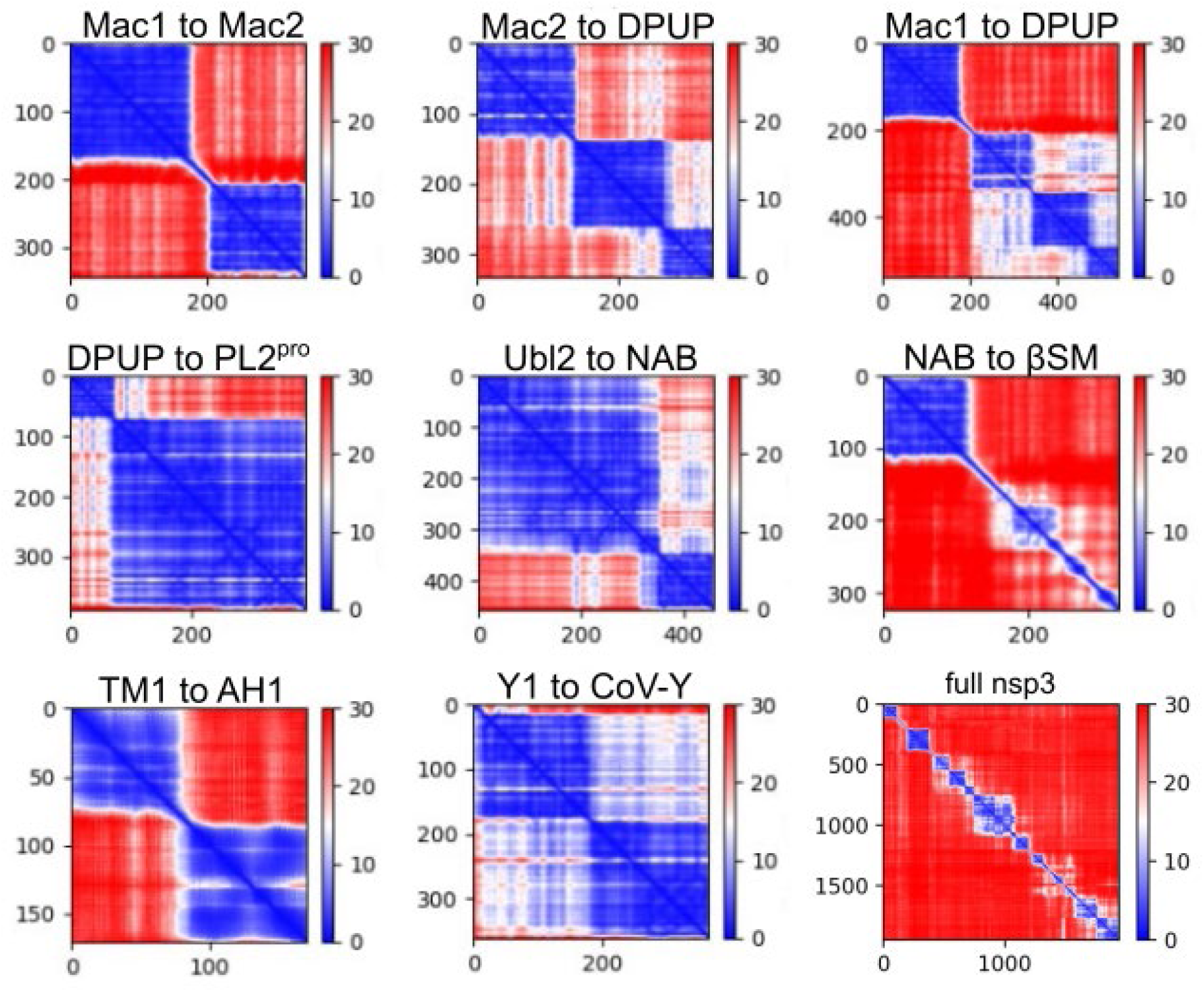
predicted Alignment Error (pAE) matrices from predicted structures which cover multiple domains from SARS-CoV-2 nsp3. Lower pAE values describe a higher confidence in the prediction of the spatial arrangement of a residue pair. Usually, blue squares in the matrix describe a folded domain, while red regions stand for an uncertain arrangement between two regions in the protein. White regions in the matrix describe an arrangement of moderate confidence between two domains. Note that the lower right matrix of the full length nsp3 prediction makes the domains and their location on nsp3 visible but shows AlphaFold2 failing to predict a confident fold of the complete protein.

### Experimental validation of the Betacoronavirus-specific linker domain (βSLD)

#### Construct design

The used construct comprises the domains Ubl2+PL2^pro^+ßSLD from SARS-CoV-2 (NCBI reference YP_009742610.1) and consists of the 345 nsp3 residues 746-1090 with the following amino acid sequence:

EVRTIKVFTTVDNINLHTQVVDMSMTYGQQFGPTYLDGADVTKIKPHNSHEGKTFYVLPNDDTLRVEAFEYYHTTDPSFLGRYMSALNHTKKWKYPQVNGLTSIKWADNNSYLATALLTLQQIELKFNPPALQDAYYRARAGEAANFCALILAYCNKTVGELGDVRETMSYLFQHANLDSCKRVLNVVCKTCGQQQTTLKGVEAVMYMGTLSYEQFKKGVQIPCTCGKQATKYLVQQESPFVMMSAPPAQYELKHGTFTCASEYTGNYQCGHYKHITSKETLYCIDGALLTKSSEYKGPITDVFYKENSYTTTIKPVTYKLDGVVCTEIDPKLDNYYKKDNSYFT

The construct contains the C111S mutation, which is commonly used in crystallization of Ubl2+PL2^pro^ [13].

#### Ramachandran analysis

The predicted structure with the sequence TTIKPVTYKLDGVVCTEIDPKLDNYYKKDNSYFT has only Thr2 as a Ramachandran outlier (analysis via MolProbity [31]). From 32 residues, 26 residues are in favoured regions, while 31 of 32 are in allowed regions. The respective plots are shown in Figure S2.

**Figure S2.**
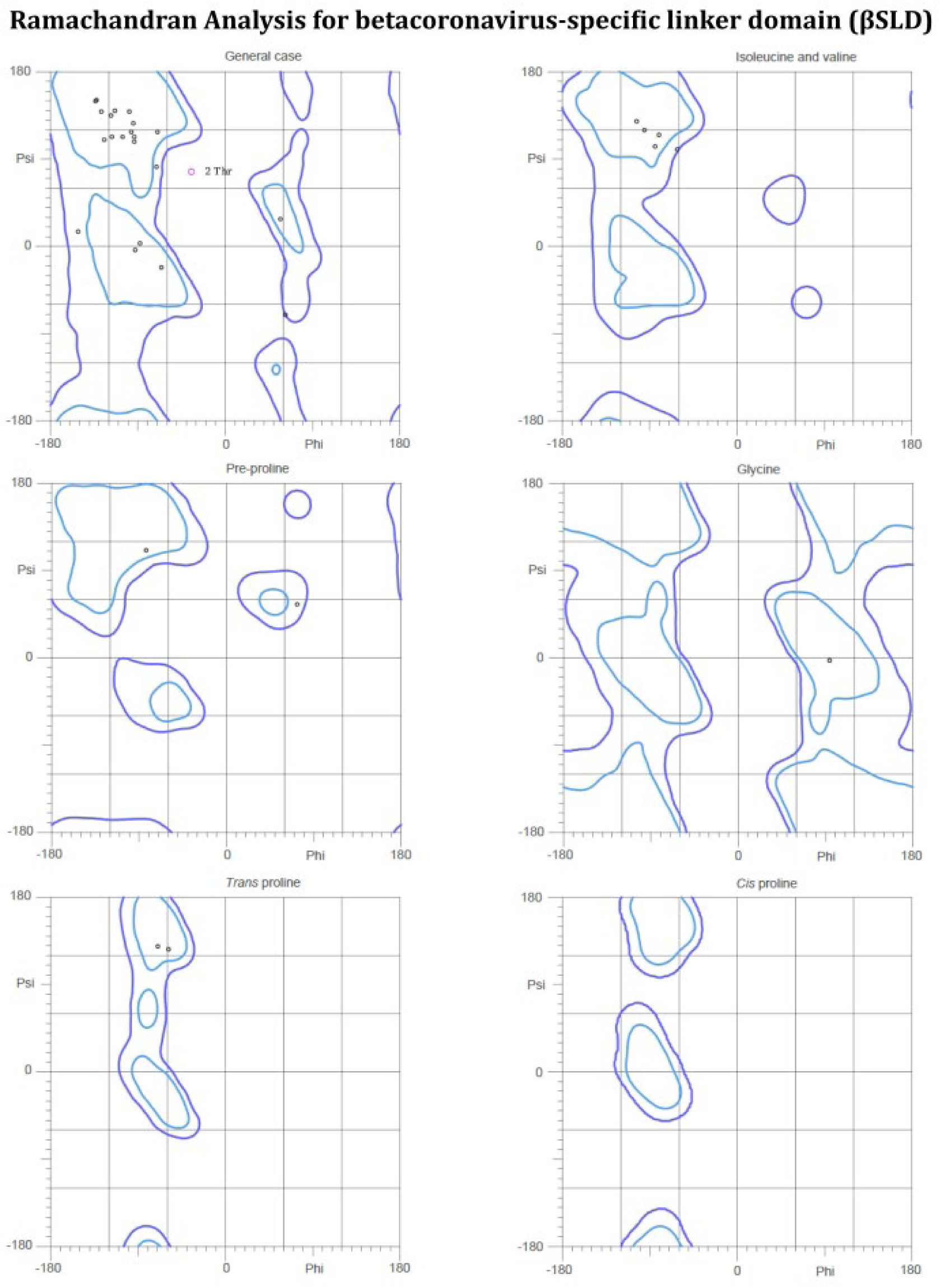
Ramachandran plots for the structure prediction of the betacoronavirus-specific linker domain.

#### Conservation of betacoronavirus-specific linker domain

Conservation of the linker domain was analysed with 57 complete genomes from *Orthocoronavirinae* (NCBI taxonomy id 2501931). This included the following NCBI accession numbers:

24 genomes from *Alphacoronavirus*:

NC_046964, NC_010437, NC_022103, NC_028811, NC_028833, NC_028814, NC_028824, NC_028752, NC_002645, NC_034972, NC_002306, NC_028806, NC_038861, NC_030292, NC_005831, NC_032730, NC_010438, NC_023760, NC_003436, NC_009988, NC_018871, NC_009657, NC_048211, NC_035191

17 genomes from *Betacoronavirus*:

NC_025217, NC_038294, NC_019843, NC_039207, NC_026011, NC_003045, NC_006213, NC_006577, NC_048217, NC_012936, NC_009020, NC_017083, NC_030886, NC_009021, NC_004718, NC_045512, NC_009019

5 genomes from *Gammacoronavirus*:

NC_010646, NC_046965, NC_048214, NC_001451, NC_010800

10 genomes from *Deltacoronavirus*:

NC_011547, NC_016996, NC_016993, NC_011550, NC_016994, NC_039208, NC_016992, NC_011549, NC_016991, NC_016995

One genome from unclassified Shrew coronavirus:

NC_046955

Due to missing annotations of non-structural proteins for most viruses, the linker domain alignment was performed mostly against pp1a and pp1ab. Most of the hits were located outside of nsp3 and AlphaFold2 predictions showed no secondary structure elements. Some of the alignments showed a similar sequence and resulted in structure predictions with a similar fold, but were located in the respective endoRNAse protein, known as nsp15 in SARS-CoV-2. These cases were NC_010646, NC_048214, and NC_010800 from *Gammacoronavirus*; all ten deltacoronaviruses, and 15 cases from *Alphacoronavirus*, namely NC_046964, NC_010437, NC_028811, NC_028833, NC_028752, NC_002645, NC_034972, NC_028806, NC_038861, NC_005831, NC_032730, NC_003436, NC_009657, NC_048211, and NC_035191.

The linker domain aligns in Deltacoronavirus endoRNAse always in the same region, which corresponds to the residues Thr275 to Lys345 in SARS-CoV-2 endoRNAse in the PDB structure 6VWW, where the case of the Night heron coronavirus HKU19 (NCBI accession number NC_016994) is depicted below:

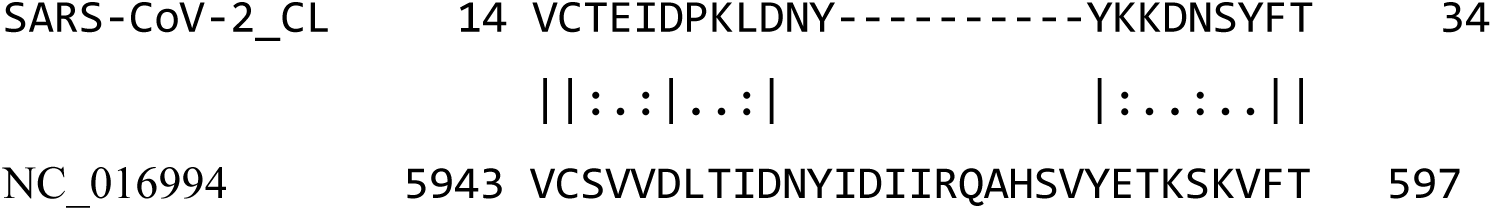

In the AlphaFold2 model, the gap-residues are predicted to form a helix, which is located right before the third β-sheet. This, however, leads in all predictions to non-compact folds with high values in the pAE matrix, where the first and the second β-sheet pairs are not connected. Therefore, we suspected the possibility that a gene duplication event occurred in an ancestor of *Betacoronavirus*, where this section of the endoRNAse translocated to nsp3 and lost this helix over time, leading to the compact fold we find in the prediction of SARS-CoV-2 βSLD. In any case, only sequence alignments with betacoronavirus sequences are located between PL2^pro^ and NAB in nsp3, indicating that this domain is specific to *Betacoronavirus*. Table S3 lists the sequence identity and similarity between SARS-CoV-2 βSLD and the 16 examined betacoronaviruses.

Sequence alignments between the linker domain of each betacoronavirus and its respective endoRNAse showed only in the case of Tylonycteris bat coronavirus HKU4 a significant hit. However, this region differs from the alignments with the other viruses.

**Table S3.**
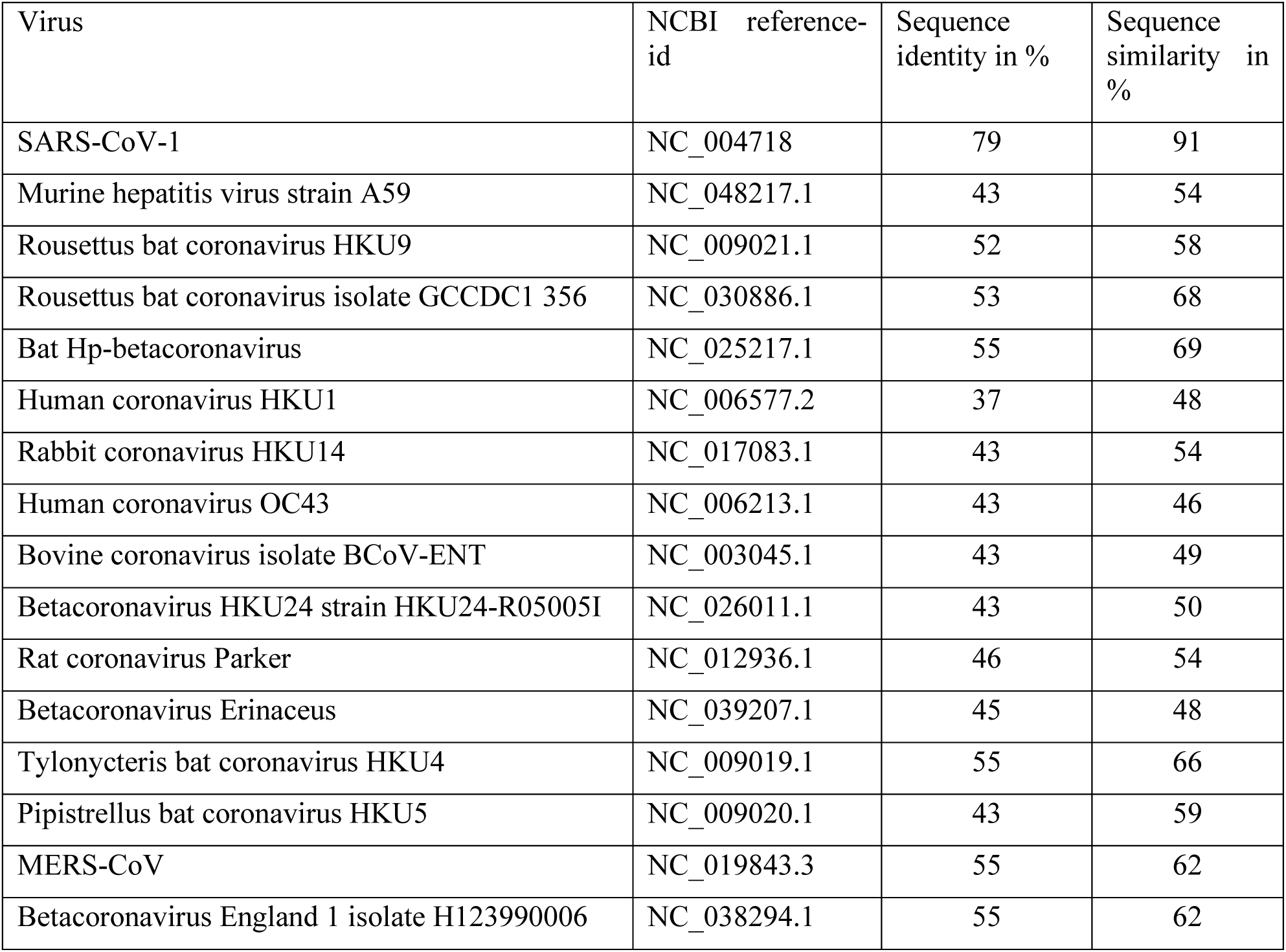
results from local pairwise sequence alignment between the sequences of several betacoronaviruses and the sequence of SARS-CoV-2 nsp3 linker domain determined by our method above. The alignment and calculations were performed via EMBOSS Water version 6.6.0 [45]. The top sequence in the alignments is always the one from SARS-CoV-2. The alignment was performed either with the sequence from an nsp3 equivalent or with the whole sequence of the pp1ab from the respective virus, if not annotated.

### Insights of the predicted hexameric assembly of Y1+CoV-Y and nsp4

#### Construct design

For validation of the Y1-hexamer, a construct of Y1 from SARS-CoV-2 (NCBI reference YP_009742610.1) with the following 161 residue sequence was used:

RATRVECTTIVNGVRRSFYVYANGGKGFCKLHNWNCVNCDTFCAGSTFISDEVARDLSLQFKRPINPTDQSSYIVDS VTVKNGSIHLYFDKAGQKTYERHSLSHFVNLDNLRANNTKGSLPINVIVFDGKSKCEESSAKSASVYYSQLMCQPIL LLDQALV

#### Conservation of Y1 in Betacoronavirus

For identification of conserved residues in *Betacoronavirus*, the same 17 betacoronaviruses were used as for the betacoronavirus-specific linker domain, listed above. A sequence of SARS-CoV-2 Y1 is found in the following lines, with a line indication each residue’s conservation status below each line of sequence:

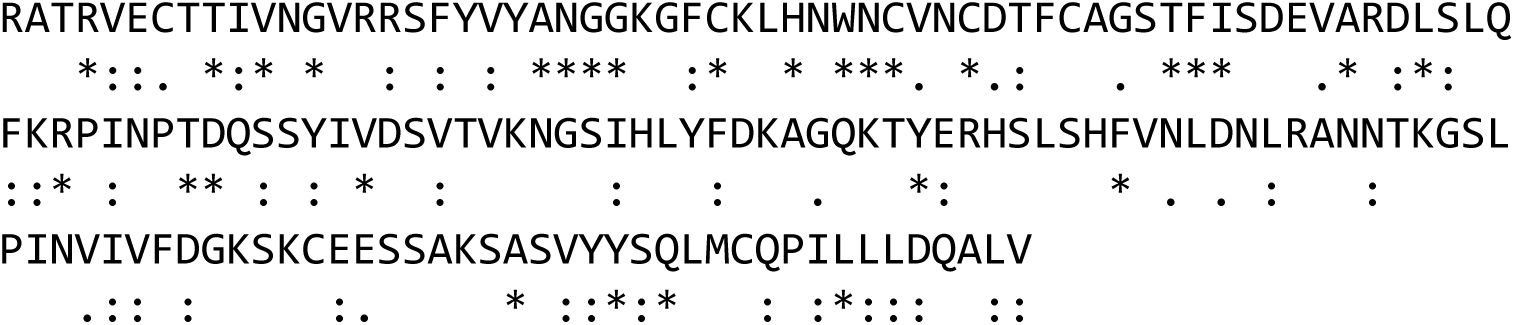

Residues marked with an asterix are conserved among all examined viruses; columns indicate high similarity of chemical properties; dots indicate low similarity of chemical properties. The 29 conserved residues with numbering based on this Y1 sequence are Arg4, Thr9, Val11, Gly13, Ala22, Asn23, Gly24, Gly25, Cys29, His32, Trp34, Asn35, Cys36, Cys39, Thr47, Phe48, Ile49, Ala54, Leu57, Arg63, Thr68, Asp69, Val75, Tyr97, Phe105, Ala141, Tyr145, Gln147, and Leu154. The following of these residues make H-bond contact to other monomers in the multimer prediction: Arg4 with Thr9; Arg4 with Asp56; Thr9 with Arg4; Gly13 with Ser17. Other residues making contact with other monomers are the non-conserved residues Ser17, Arg44, Asp56, Leu120, Ser131, Ser136, Lys139, and Leu148. In total, eleven residues make monomer-contact in the multimer prediction of SARS-CoV-2 Y1.

#### Conservation of Y1 in Coronaviridae

For this analysis, we used one or two representatives of one genus, given here with the NCBI reference id. These were NP_073549.1 from *Alphacoronavirus*, NP_066134.1 and YP_001876435.1 from *Gammacoronavirus*, YP_002308496.1 and YP_002308505.1 from *Deltacoronavirus*, and NC_046955.1 from the unclassified Shrew coronavirus. Sequence identities range from 25.95% to 39.10% and RMSD values in comparison of structure predictions to the prediction of SARS-CoV-2 Y1 ranged from 0.62 Å to 1.28 Å. While not all available viruses were examined, the few representatives show high similarity, indicating a conservation in *Coronaviridae*.

#### Conservation of Y1 in Nidovirales

For related viruses outside of *Coronaviridae*, we compared the sequence to viruses from *Pitovirinae*, *Letovirinae*, *Torovirinae, Roniviridae*, *Mesoniviridae*, *Arteriviridae* and *Piscanivirinae*, which are all mentioned in Neumann et al. [14]. The sequence identities ranged from 17.95% to 24.36%, whereas most cases showed identities below 20%. Structure prediction showed no similar structures, with RMSD values ranging from 5.96 Å to 15.8 Å. The gill-associated virus (*Roniviridae*, NCBI reference: YP_001661453.1) has a sequence identity of 21.7%, and the prediction shows a partially confident fold, but even after extending the sequence in multiple iterations, none of the predictions resembled the coronaviral Y1. The same observation was made with sequences from *Mesoniviridae*, *Torovirinae*, *Piscanivirinae*, and the unclassified bovine nidovirus (NCBI reference: YP_009142788.1).

#### Multimer prediction of SARS-CoV-2 and MHV Y1+CoV-Y

The multimer prediction of Y1+CoV-Y shows higher pLDDT and better pAE values when predicted for MHV compared to SARS-CoV-2 (Figure S3). Both structures are generally similar, but the CoV-Y to Y1 arrangement deviates, leading to an RMSD of 2.5 Å between SARS-CoV-2 and MHV structure.

**Figure S3.**
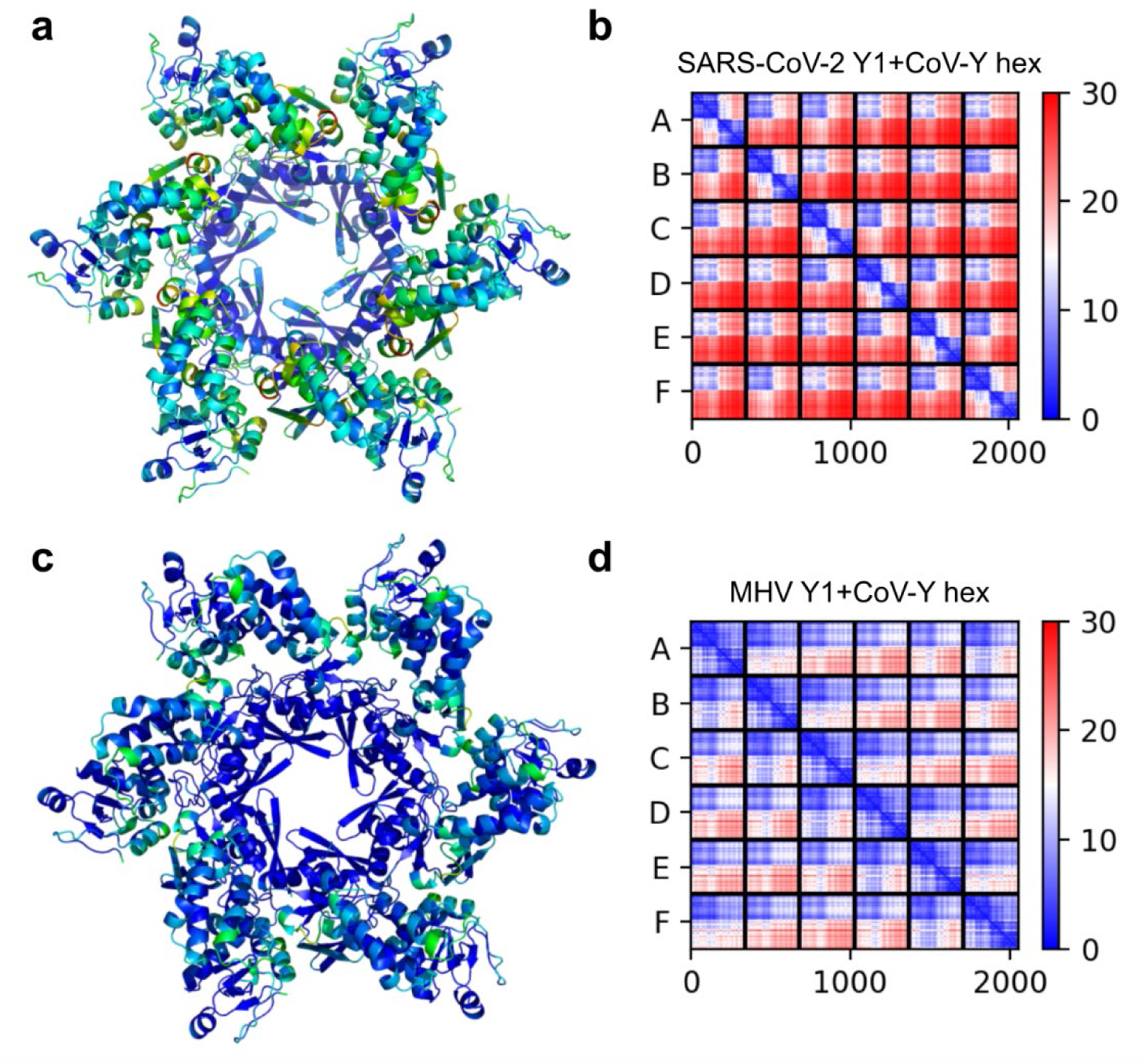
Comparison of SARS-CoV-2 and MHV hexamer prediction of Y1+CoV-Y. a: hexamer prediction of SARS-CoV-2 Y1+CoV-Y. b: pAE matrix for structure shown in (a). c: hexamer prediction of MHV Y1+CoV-Y. d: pAE matrix for structure shown in (c). (a) and (c) are colored according to pLDDT, with values above 90 shown in deep blue and values below 50 shown in red, with other colors interpolated in between. Y1 domains in (c) are arranged well according to pAE values in (d). CoV-Y is only arranged well to the Y1 of the same monomer and in some cases also well to Y1 of nearby monomers.

#### Ectodomain

During identification of ordered and disordered regions via AlphaFold2, we also identified the folded core of the ectodomain, the only nsp3 domain located at the lumenal side of the membrane. It is not experimentally solved and according to pLDDT and pAE values, only a fraction of the lumenal residues folds into a compact domain.

Between the two sarbecoviruses and MHV, the folding part of the ectodomain shows a high sequence similarity. However, the structure prediction shows two distinct folds: one for the sarbecoviruses, with three short α-helices interconnected by two disulfide bonds; and one for MHV, with only one α-helix, two β-sheets and only one disulfide bond (Figure S4). However, the four cysteines are present in all three viruses. The structure is of relevance to understanding nsp3 since the ectodomain was shown to interact with a lumenal domain of nsp4, which leads to the formation of the replication organelles [6]. If the predicted structures can be validated experimentally, it would be interesting to examine the mutations which would lead to the different folds and how this affects the interaction with nsp4.

**Figure S4.**
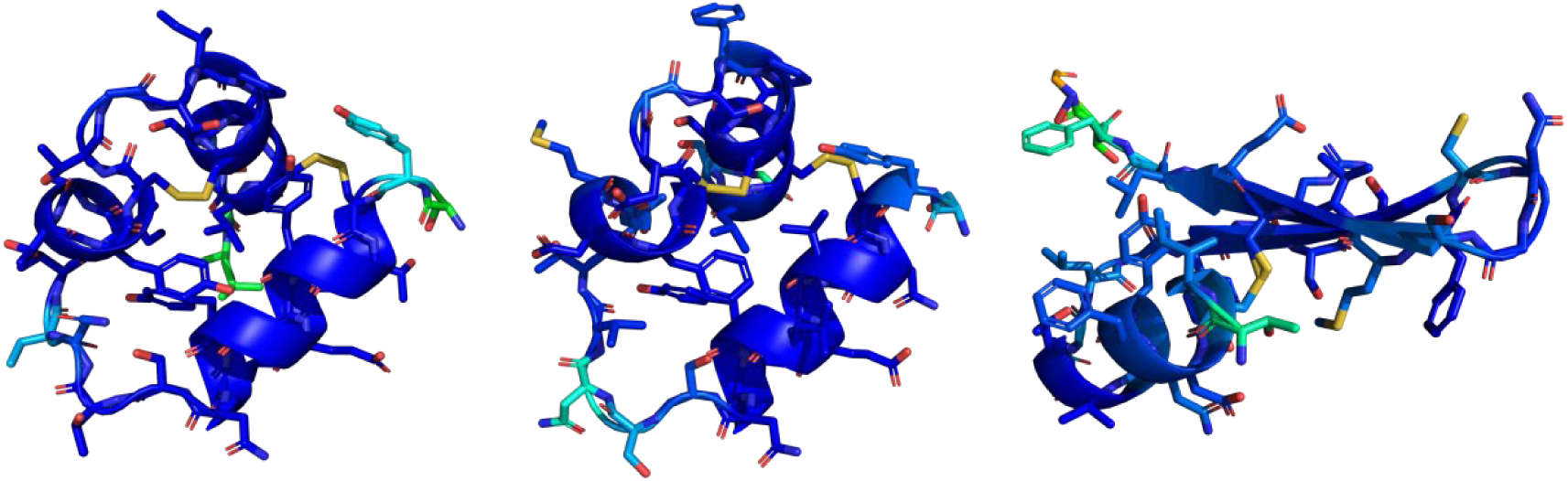
Comparison structure predictions of the ectodomain between SARS-CoV-2 (left), SARS-CoV-1 (center), and MHV (right), colored according to pLDDT with values below 50 in red and values of 90 in deep blue, with other colors interpolated in between. Non-carbon atoms are colored by element. Note that two disulfide bonds are present in the sarbecoviruses, while only one is present in MHV and that MHV ectodomain shows a different fold containing β-sheets.

#### Domains and multimer prediction of Nsp4

Previous research revealed nsp4 to be part of the hexameric pore complex [7] and identified residues, which interact with the ectodomain of nsp3 [6]. The domains of nsp4 were analysed analogously to nsp3 with the support of transmembrane domain prediction and AlphaFold2, which reveals a large nsp4 ectodomain surround by two transmembrane domains, as well as a small C-terminal domain and two additional transmembrane helices. The domain ranges for nsp4 are listed in table S4.

A multimer-prediction of the nsp4 ectodomain results in a hexameric pore with an inner diameter of 2-3 nm, which matches with experimental measurements of the pore [7]. However, the pAE matrix shows weak confidence in the arrangement of monomers and the orientation of the termini, which are attached to transmembrane helices, contradict experimentally proven membrane topology [5].

**Table S4.** domain ranges of nsp4 for both sarbecoviruses and MHV. Ranges of transmembrane domains are predicted by TMHMM 2.0 [27]. The N-terminal tail and all linkers show low pLDDT values in AlphaFold2 predictions and are classified as likely disordered.

| Nsp4 Domain | SARS-CoV-2 ranges | SARS-CoV-1 ranges | MV ranges |
| --- | --- | --- | --- |
| N-terminal tail | 1-12 | 1-12 | 1-6 |
| Transmembrane domain 1 | 13-32 | 13-35 | 7-29 |
| Nsp4-Ectodomain | 33-279 | 36-279 | 30-278 |
| Transmembrane domain 2 | 280-302 | 280-302 | 279-301 |
| Linker TM2-TM3 | 303-313 | 303-314 | 302-312 |
| Transmembrane domain 3 | 314-348 | 315-337 | 313-335 |
| Linker TM3-TM4 | 349-362 | 338-364 | 336-370 |
| Transmembrane domain 4 | 363-385 | 365-387 | 371-393 |
| C-terminal folded domain | 386-500 | 388-500 | 394-496 |

